# Engineering the cyanobacterial ATP-driven BCT1 bicarbonate transporter for functional targeting to C_3_ plant chloroplasts

**DOI:** 10.1101/2024.02.14.580295

**Authors:** Sarah Rottet, Loraine M. Rourke, Isaiah C.M. Pabuayon, Su Yin Phua, Suyan Yee, Hiruni N. Weerasooriya, Xiaozhuo Wang, Himanshu S. Mehra, Nghiem D. Nguyen, Benedict M. Long, James V. Moroney, G. Dean Price

**Author notes:** Co-first author (Sarah Rottet, Loraine M. Rourke).

## Abstract

The ATP-driven bicarbonate transporter 1 (BCT1), a four-component complex in the cyanobacterial CO_2_-concentrating mechanism, could enhance photosynthetic CO_2_ assimilation in plant chloroplasts. However, directing its subunits (CmpA, CmpB, CmpC and CmpD) to three chloroplast sub-compartments is highly complex. Investigating BCT1 integration into *Nicotiana benthamiana* chloroplasts revealed promising targeting strategies using transit peptides from the intermembrane space protein Tic22 for correct CmpA targeting, while the transit peptide of the chloroplastic ABCD2 transporter effectively targeted CmpB to the inner envelope membrane. CmpC and CmpD were targeted to the stroma by RecA and recruited to the inner envelope membrane by CmpB. Despite successful targeting, expression of this complex in CO_2_-dependent *Escherichia coli* failed to demonstrate bicarbonate uptake. We then used rational design and directed evolution to generate new BCT1 forms that were constitutively active. Several mutants were recovered, including a CmpCD fusion. Selected mutants were further characterized and stably expressed in *Arabidopsis thaliana,* but the transformed plants did not have higher carbon assimilation rates or decreased CO_2_ compensation points in mature leaves. While further analysis is required, this directed evolution and heterologous testing approach presents potential for iterative modification and assessment of CO_2_-concentrating mechanism components to improve plant photosynthesis.

**Highlight:** We describe the directed evolution and rational design of a cyanobacterial four-component bicarbonate transporter and the localization of its subunits to various chloroplast sub-compartments for improving C_3_ plant photosynthesis.

## Introduction

A crop improvement approach of ongoing global interest is the utilisation of cyanobacterial and algal CO_2_-concentrating mechanisms (CCMs) to enhance photosynthetic performance through improved carbon fixation (Price *et al*., 2013; Long *et al*., 2016; Hennacy and Jonikas, 2020; Nguyen *et al*., 2024). Carboxylation of ribulose-1,5-bisphosphate by the bifunctional enzyme ribulose-1,5-bisphosphate carboxylase/oxygenase (Rubisco) is a major limitation to efficient carbon acquisition by crops (Long *et al*., 2015). Cyanobacterial and algal CCMs, however, have evolved to actively accumulate bicarbonate (HCO_3-_) within cellular compartments to supply high CO_2_ concentrations to fast Rubisco enzymes for highly efficient carbon acquisition (Rae *et al*., 2017). A number of strategies exist for the creation of a functional CCM in C_3_ crop plants (Moroney *et al*., 2023), but crucial to all of these is a requirement to increase HCO_3-_ concentration in the chloroplast stroma to supply either the native Rubisco, or one that has evolved in a CCM, so that the CO_2_ fixation reaction is optimized (Price *et al*., 2011; Rottet *et al*., 2021).

Cyanobacterial and algal CCMs utilise a suite of dedicated bicarbonate transporters that consume cellular energy to elevate HCO_3-_ ion concentrations inside cellular membranes (Rae *et al*., 2017; Rottet *et al*., 2021), to levels up to 1,000-fold higher than the external environment (Price *et al*., 2008). Since passive diffusion of HCO_3-_ across membranes is very slow compared to CO_2_ (Tolleter *et al*., 2017), a key to CCM function is that active bicarbonate pumping leads to the successful elevation of HCO_3-_ inside cells. Then, specific carbonic anhydrase (CA) enzymes located with Rubisco interconvert the accumulated HCO_3-_ to CO_2_, enabling a localized elevation of CO_2_ for use by Rubisco (Moroney *et al*., 2023). Within crop-CCM strategies, the successful elevation of chloroplastic HCO_3-_ concentrations via bicarbonate transporters alone is expected to provide increased photosynthetic output through provision of a net increase in CO_2_ supply to Rubisco (Price, 2011; McGrath and Long, 2014; Wu *et al*., 2023).

To date, efforts to successfully express and deliver functional bicarbonate transporters to the correct location in plants have highlighted complexity with respect to protein targeting and function in crop systems (Pengelly *et al*., 2014; Atkinson *et al*., 2016; Rolland *et al*., 2016; Uehara *et al*., 2016, 2020; Nölke *et al*., 2019; Förster *et al*., 2023). Those studies predominantly addressed the use of relatively simple, single or dual gene bicarbonate pump systems (e.g. SbtA/B, BicA, LCIA, HLA3), as opposed to the more complex higher-order bicarbonate pumps and CO_2_-to-bicarbonate conversion complexes found in native CCMs (Rottet *et al*., 2021). Despite these complexities, some higher-order bicarbonate pumps present desirable characteristics for HCO_3-_ accumulation in the chloroplast stroma such as energization and no ion co-transport dependencies (Rottet *et al*., 2021).

Here we address the potential to make use of a relatively complex bicarbonate pump system, bicarbonate transporter 1 (BCT1), in the engineering of crop chloroplast CCMs. BCT1 is an ideal candidate for HCO_3-_ accumulation in the chloroplast stroma, owing to its high affinity for bicarbonate, its ability to transport HCO_3-_ against a concentration gradient, and because it is energized by ATP hydrolysis. In cyanobacteria, BCT1 is a low-inorganic carbon (Ci)-inducible ATP-binding cassette (ABC) transporter encoded by the *cmpABCD* operon under the control of the transcriptional regulator CmpR (Omata *et al*., 1999*b*, 2001; Nishimura *et al*., 2008; Pan *et al*., 2016). The operon gives rise to the expression of four protein components; CmpA, CmpB, CmpC and CmpD which occupy different locations associated with the cyanobacterial plasma membrane (*Figure 1A*). The *cmpABCD* operon is found in both α- and β-cyanobacterial species (Rae *et al*., 2011; Sandrini *et al*., 2014; Cabello-Yeves *et al*., 2022), and therefore a ubiquitous element in cyanobacterial CCMs.

**Figure 1.**
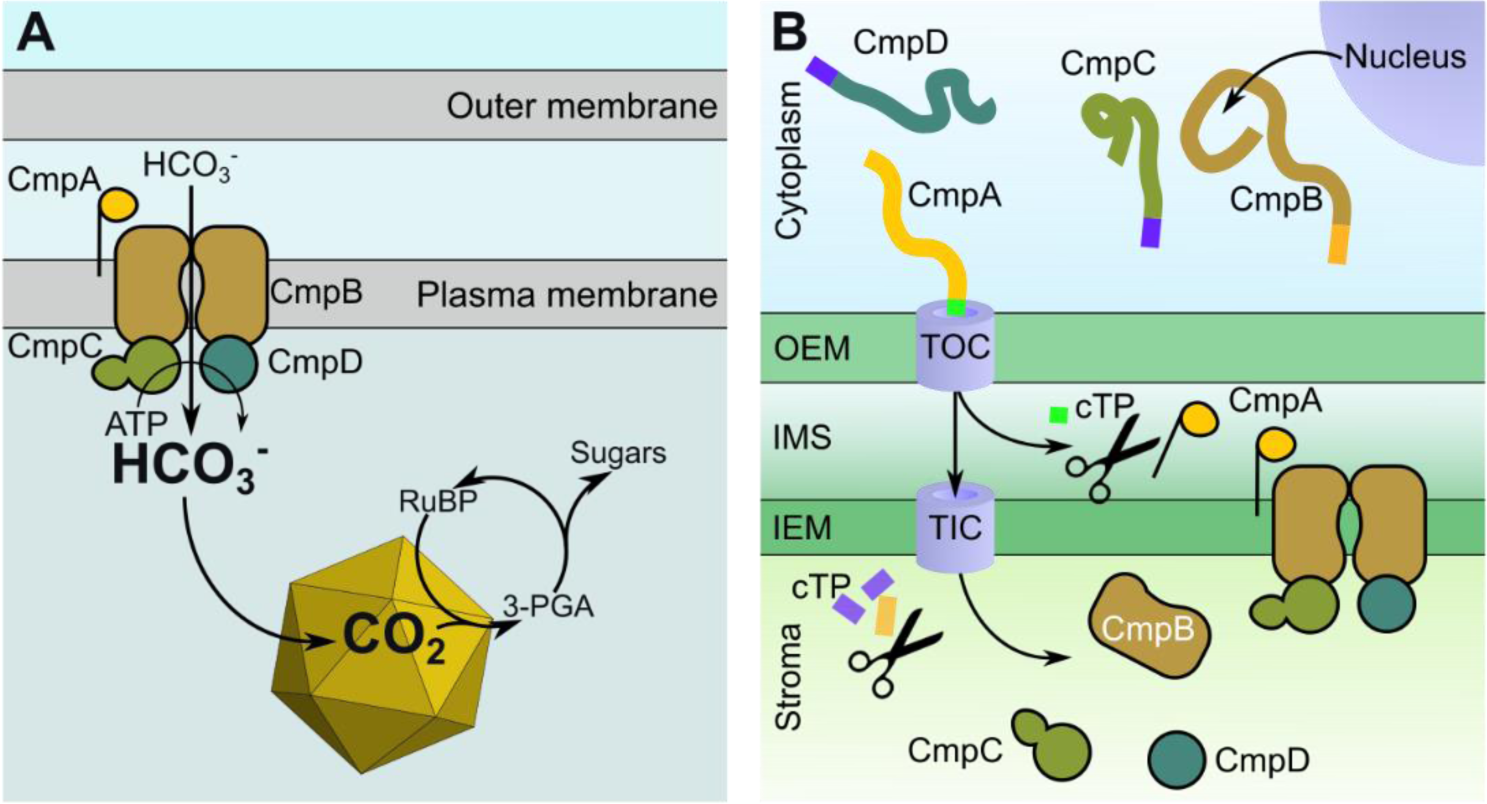
Structure of BCT1 and strategy for its installation in the chloroplast envelopes. **(A)** In cyanobacteria, BCT1 transports HCO_3-_ across the plasma membrane. Firstly, HCO_3-_ is captured by the substrate-binding protein CmpA and delivered to the membrane protein CmpB. HCO_3-_ travels across the plasma membrane through the channel formed by a homodimer of the protein CmpB. CmpC and CmpD are nucleotide-binding proteins or ATPases which sit inside the cyanobacterial cell and hydrolyse ATP to provide the energy for the transport of HCO_3-_ across the plasma membrane. Once in the cell, HCO_3-_ diffuses into the carboxysome where it is converted into CO_2_ by a carbonic anhydrase. **(B)** The strategy for the installation of the cyanobacterial BCT1 complex in the chloroplast envelope is based on nucleus-encoded CmpA, CmpB, CmpC and CmpD. Each protein is individually targeted to the appropriate chloroplast sub-compartment using three different chloroplast transit peptides (cTP). CmpA is targeted to the intermembrane space (IMS). CmpB is sent to the inner envelope membrane (IEM), while CmpC and CmpD are targeted to the stroma. OEM, outer envelope membrane; TOC, translocon at the outer envelope membrane of chloroplasts; TIC, translocon at the inner envelope membrane of chloroplasts.

BCT1 is a high affinity transporter, exhibiting an apparent *K_m_* of 15 µM for HCO_3-_ (Omata *et al*., 1999*b*), and is a multi-subunit ABC transporter, closely related to the nitrate transporter NrtABCD (Omata, 1995; Klanchui *et al*., 2017). The substrate-binding protein (SBP) component, CmpA, binds HCO_3-_ with high affinity (*K_d_* = 5 µM) and transfers it to the membrane transport complex (Maeda *et al*., 2000). The first 28 N-terminal residues of CmpA form a lipoprotein signal peptide, which, when removed, results in a functional soluble protein in *Escherichia coli* (Maeda *et al*., 2000). The signal peptidase II recognises the cleavage site ^26^LKGC^29^, which, following cleavage and removal of the lipoprotein, creates a covalent bond between CmpA and lipids via Cys^29^ (Maeda and Omata, 1997; Tjalsma *et al*., 1999). CmpB is the transmembrane domain (TMD) component of BCT1 and is likely to form a homodimer that functions as the channel for the transport of HCO_3-_ across the plasma membrane (Omata *et al*., 2002). Finally, the nucleotide-binding domain (NBD) proteins, CmpC and CmpD, are likely to form a heterodimer that hydrolyses ATP to power the transport of HCO_3-_ (Omata *et al*., 1999*a*; Smith *et al*., 2002). In cyanobacteria, both sit on the cytoplasmic side of the plasma membrane (*Figure 1A*). CmpD is a canonical NBD containing highly conserved ATP binding motifs (i.e. Walker A, Walker B, ABC signature; Schneider and Hunke, 1998). In contrast, CmpC is a non-canonical NBD harboring an additional C-terminal domain that is 50% similar to NrtA, homologous to CmpA, and thought to act as a solute-binding regulatory domain.

The engineering complexity of constructing a functional form of BCT1 in a crop plant chloroplast is evidenced by the requirement for each protein component of the BCT1 complex to be targeted to a specific sub-compartment of the chloroplast. Given the limited applicability of plastome transformation technologies across diverse crop species (Hanson *et al*., 2013), we here use a nuclear transformation approach, which has broader applicability (*Figure 1B*). Our previous work demonstrated that unmodified BCT1 had no bicarbonate uptake activity when expressed in *E. coli* (Du *et al*., 2014). This highlights the potential requirement for regulatory systems that exist in cyanobacteria to modify BCT1 function, such as post-translational phosphorylation (Spät *et al*., 2021), suggesting the requirement of other factors external to the complex itself in order to control function. Moreover, CmpC regulatory domain function is not yet fully understood., potentially due to the absence of native regulatory mechanisms.

Here, we investigated strategies for targeting *Synechococcus sp*. PCC7942 BCT1 subunits to plant chloroplast locations and used mutagenesis to obtain variants with activation independent of unidentified control mechanisms. A synthetic biology approach that combined chloroplast sub-compartment targeting peptides with fluorescent reporter proteins was used to identify the best targeting systems for each BCT1 component. We also employed a directed evolution approach in a specialized *E. coli* strain that lacks CA (hereafter CA-free) and requires high levels of CO_2_ for growth (Du *et al*., 2014; Desmarais *et al*., 2019; Förster *et al*., 2023). We were therefore able to control the function of a stand-alone BCT1 complex and eliminate regulatory requirements absent in heterologous systems. This resulted in the generation of constitutively active forms of BCT1 in *E. coli*. However, the expression of BCT1 in Arabidopsis did not result in the expected elevation in CO_2_ supply to Rubisco. Although the tested BCT1 constructs did not exhibit functionality in Arabidopsis at this stage, our work has established a framework to assess correct protein targeting in *N. benthamiana*, activity in *E. coli*, and eventual functionality in plants. We have developed tools for assessing bicarbonate uptake activity *in vivo* in both *E. coli* and Arabidopsis. Moving forward, this process is likely to be iterative, with the next steps involving the evaluation of constructs generated through directed evolution to ascertain their targeting efficiency and expression levels in Arabidopsis.

## Results

### Individual targeting of BCT1 components to the chloroplast

To determine the optimal route for installing BCT1 subunits to the correct location in chloroplasts, we employed a transient expression approach in *Nicotiana benthamiana* combined with fluorescent reporter constructs and confocal microscopy. Targeting foreign proteins to specific chloroplast sub-compartments is a significant engineering challenge as there are at least six sub-compartments (i.e. outer envelope membrane [OEM], intermembrane space [IMS], inner envelope membrane [IEM], stroma, thylakoid membrane, and thylakoid lumen; Rolland *et al*., 2017). Specifically, we targeted nucleus-encoded CmpA, CmpB, CmpC, and CmpD individually to the chloroplast IMS, IEM, or stroma using a variety of chloroplast transit peptides (cTP; *Figure 1B*).

To date, the targeting of only two IMS proteins, Tic22 and MGD1, have been studied (Kouranov *et al*., 1999; Vojta *et al*., 2007; Chuang *et al*., 2021). While *At*MGD1 cTP targeted CmpA to the stroma, Tic22 isoforms from *Arabidopsis thaliana* and *Pisum sativum* proved more successful in targeting CmpA to the IMS (*Supplementary Figure S1*). Notably, the first 64 residues of the protein *At*Tic22-IV targeted CmpA to the IMS of *N. benthamiana* (*At*Tic22-IV_64_-CmpA, *Figure 2*).

**Figure 2.**
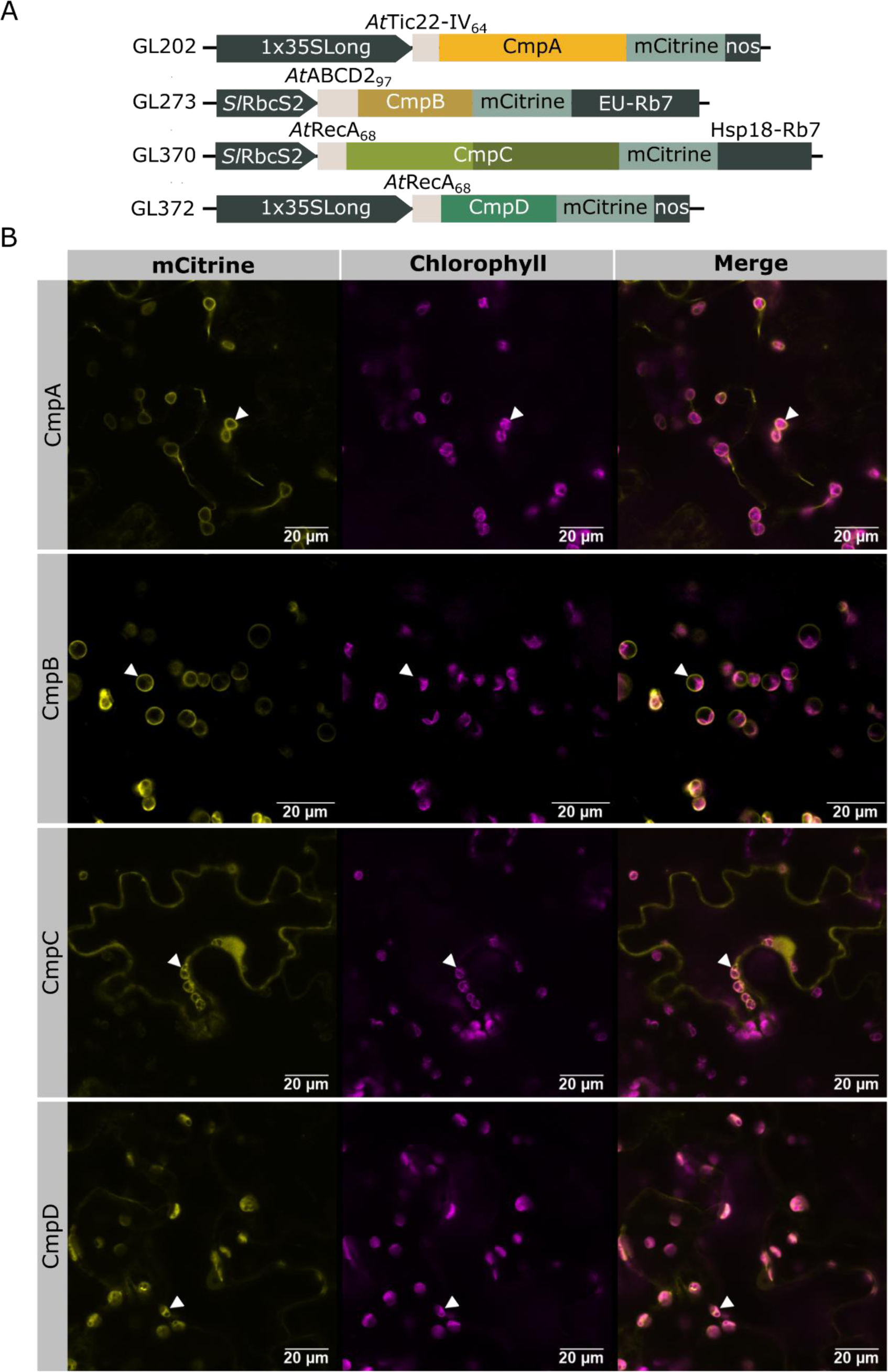
Individual targeting of CmpA, CmpB, CmpC, and CmpD to Nicotiana benthamiana chloroplasts. **(A)** Schematic of the genetic constructs used in this figure. The chloroplast transit peptides (cTPs) originate from *Arabidopsis thaliana* (At). The proteins used are *At*Tic22-IV (At4g33350, GL202), *At*ABCD2 (At1g54350, GL273), and *At*RecA (At1g79050, GL370, GL372). The length of the cTPs are shown as the number of residues in subscript. BCT1 genes are coloured as in *Figure 1*. CmpC NBD and regulatory domain are shown in light and dark green respectively. **(B)** Confocal microscopy images of *N. benthamiana* leaf surfaces transiently expressing BCT1 proteins fused with mCitrine. CmpA localized at the chloroplast intermembrane space (arrow head), CmpB at the inner envelope membrane (arrow head), and CmpD in the stroma (arrow head). CmpC localized in the stroma (arrow head) and in the cytosol.

To target CmpB to the IEM, ABC transporters predicted to localize to the IEM were identified from chloroplast proteomes (Ferro *et al*., 2010; Simm *et al*., 2013; Bouchnak *et al*., 2019). The targeting efficiency of a subset of leader sequences from ABC transporters (*At*TAP1, *At*ABCD2, *At*ABCG7) and other candidates (e.g. *At* PLGG1_92_; Rolland et al., 2016) were assessed (*Supplementary Figure S2*). We found that the first 97 residues of *At*ABCD2 transporter effectively targeted CmpB to the IEM (*At*ABCD2_97_-CmpB, *Figure 2*), while some targeting sequences (i.e. *At*ABCG7) completely failed to deliver CmpB to the chloroplast (*Supplementary Figure S2*).

In cyanobacteria, CmpC and CmpD are cytoplasmic NBD components of the BCT1 complex and are expected to bind transiently to their membrane anchor, CmpB. As a result, in a chloroplastic CCM, targeting of CmpC and CmpD to the IEM is unnecessary. Instead, we attempted to target them to the chloroplast stroma. To achieve this, we employed the well-established stromal targeting sequence from *At*RecA (Köhler *et al*., 1997). While it efficiently targeted CmpD to the stroma, CmpC targeting was effective but less efficient, also being detected in the cytosol after 3-days post-infiltration (*At*RecA_68_-CmpD and *At*RecA_68_-CmpC, *Figure 2*).

### Recruitment of CmpC and CmpD to the inner envelope membrane by CmpB

To determine whether CmpB is properly oriented in the membrane to interact with its stromal NBDs, a strategy involving co-expression of individual NBD with CmpB in *N. benthamiana* was employed. CmpC and CmpD were tagged with fluorescent reporters, while CmpB carried a small non-fluorescent label (HA-H_6_) to reduce potential interference (*Figure 3A*). Confocal microscopy was used to track NBD localization and detect a shift from the stroma to the IEM.

**Figure 3.**
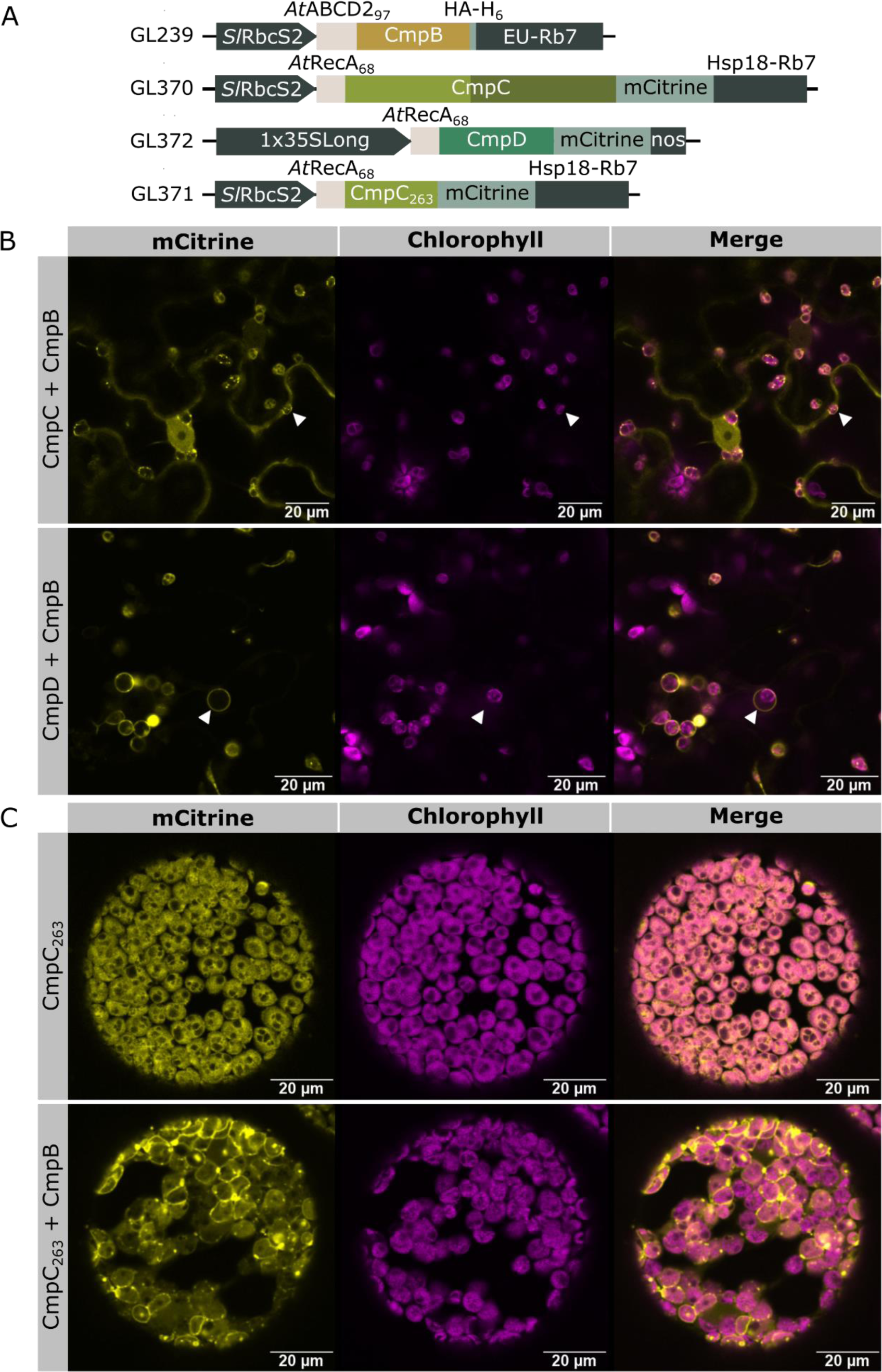
Combinatorial targeting of CmpC or CmpD with CmpB to the chloroplasts of Nicotiana benthamiana. **(A)** Schematic of the genetic constructs used in this figure. The chloroplast transit peptides (cTPs) originated from *At*ABCD2 (At1g54350, GL239), and *At*RecA (At1g79050, GL370-372). The length of the cTPs are shown as the number of residues in subscript. CmpB (GL239) is tagged with the non-fluorescent HA-H_6_ epitope, while CmpC (GL370), CmpC_263_ (GL371) and CmpD (GL372) are fused with mCitrine. **(B)** Confocal microscopy images of *N. benthamiana* leaf surfaces transiently expressing a combination of two BCT1 proteins. When CmpC was co-expressed with CmpB (row 1), CmpC mostly remained in the cytosol but seemed to also localize at the IEM (arrow head). When CmpD and CmpB were co-expressed (row 2), CmpD clearly localized at the IEM (arrow head). **(C)** Confocal microscopy images of *N. benthamiana* protoplasts transiently expressing a truncated form of CmpC that lacks the regulatory domain (CmpC_263_). Individual targeting of CmpC_263_ (row 1) resulted in a stromal localization pattern, while co-expression with CmpB (row 2) led to the relocalization of CmpC_263_ to the IEM.

While *At*RecA_68_-CmpD (GL372) alone was targeted to the stroma, it was successfully recruited to the IEM when co-expressed with *At*ABCD2_97_-CmpB (GL239; *Figure 3B*). We could not obtain conclusive evidence of *At*RecA_68_-CmpC (GL370) relocalization to the IEM when co-expressed with *At*ABCD2_97_-CmpB, likely due to its slow delivery to the chloroplast and relative accumulation in the cytosol (*Figure 3B*). However, the removal of the regulatory domain in CmpC (*At*RecA_68_-CmpC_263_, GL371) allowed for cleaner targeting to the stroma and obvious recruitment to the IEM by *At*ABCD2_97_-CmpB (*Figure 3C*). Successful recruitment of the two NBDs suggests that *At*ABCD2_97_-CmpB not only sits in the chloroplast IEM but is also in the correct orientation to allow appropriate protein:protein interactions with stromal CmpC and CmpD.

Considering the limited understanding of membrane protein orientation determinants, we utilized our system to explore the influence of various targeting sequences on the orientation of CmpB in the membrane. Although some targeting sequences were less efficient in delivering CmpB to the IEM, they did not affect its orientation. All seven tested targeting sequences for CmpB triggered the relocalization of *At*RecA_54_-CmpC_263_ (GL199) to the IEM (*Supplementary Figure S3*). In contrast, the control construct lacking a targeting sequence for CmpB (GL234, no SP) did not induce the shift of CmpC_263_ from the stroma to the IEM (*Supplementary Figure S3*).

### Generation of active BCT1 mutants by rational design

Previous results showed that unmodified BCT1 is inactive in *E. coli* (Du *et al*., 2014). To address this, we initially removed putative regulatory requirements of BCT1 by rational design. For this purpose, we used a Loop Assembly (Pollak *et al*., 2019) approach, enabling high throughput design and construction of flexible linkers, point mutations, and domain deletions (*Figure 4*). Since we hypothesized the lack of BCT1 function in heterologous systems may be due to the absence of regulatory mechanisms present in cyanobacteria, an obvious rational design approach was to remove the regulatory domain of CmpC (*Figure 4C*). CmpC is a 663-residue protein, of which only 263 residues fold into a canonical NBD (*Supplementary Figure S4*). The additional C-terminal domain is thought to be involved in BCT1 regulation (Omata *et al*., 2002; Koropatkin *et al*., 2006). We therefore generated a construct that only encoded the first 263 residues of CmpC, namely CmpC_263_.

**Figure 4.**
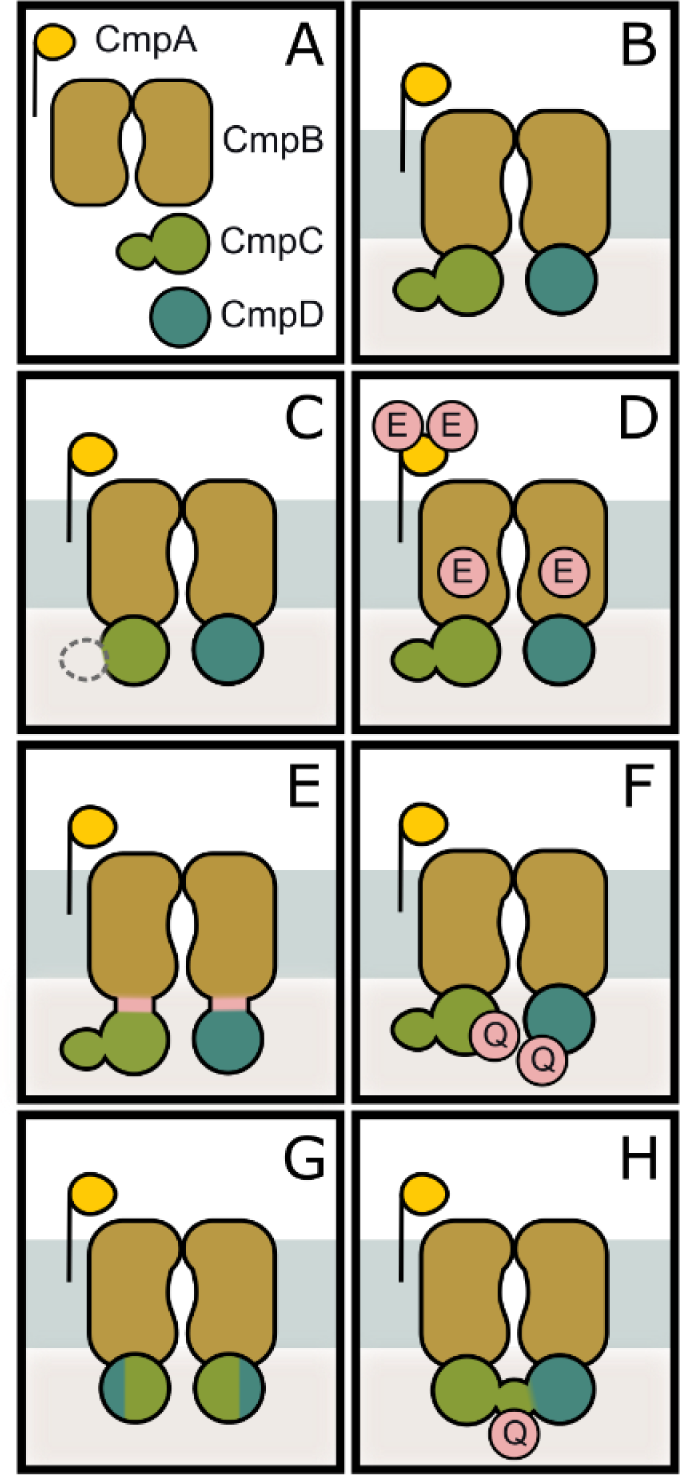
BCT1 mutants obtained by rational design and directed evolution. Schematic representation of BCT1 mutants generated by rational design **(C-F)** and directed evolution **(G-H)**. **(A)** BCT1 subunits are CmpA (gold), CmpB (brown), CmpC (green), and CmpD (teal). **(B)** Unmodified. **(C) Without** regulatory domain using CmpC_263_. **(D)** Phosphorylation mimics with CmpA^S107E, T126E^ and CmpB^T3E^. **(E)** Translational fusions of CmpBC and CmpBD (reflecting a half-transporter design, Ford *et al*., 2019). **(F)** ATP hydrolysis deficient with CmpC^E164Q^ and CmpD^E179Q^. **(G)** CmpCD chimera. **(H)** CmpCD fusion with CmpC^H409Q^. Point mutations are shown as red circles with the new residue as single letter code.

We also generated point mutations in CmpA and CmpB to mimic potential phosphorylation events found in *Synechocystis* sp. PCC6803 (CmpA: S110, T129; and CmpB: T3; Spät *et al*., 2021). We identified the corresponding residues in *Synechococcus elongatus* PCC7942 and mimicked phosphorylation by serine/threonine-to-glutamic acid substitutions in CmpA^S107E, T126E^ and CmpB^T3E^ (*Figure 4D*).

In prokaryotes, the two TMDs and two NBDs of ABC transporters are often encoded by separate genes, while in eukaryotes, these domains are typically connected by linker region(s) to form so-called ‘full-‘ or ‘half-transporters’ (Theodoulou and Kerr, 2015; Ford *et al*., 2019). Half-transporter fusions of CmpB with CmpC (hereafter CmpBC) and CmpB with CmpD (hereafter CmpBD) were generated using flexible linkers of approx. 40 residues (*Figure 4E*). This should ensure domain assembly when expressed in more complex heterologous systems and reduce targeting complexity (Ford *et al*., 2019).

In ABC transporters, it is accepted that ATP hydrolysis is carried out by the NBDs and that a glutamate-to-glutamine substitution in the conserved Walker B motif causes ATP hydrolysis deficiency (Orelle *et al*., 2003). A putatively inactive BCT1 mutant was created as a negative control (*Figure 4F*) by mutating the catalytic glutamate in both CmpC^E164Q^ and CmpD^E179Q^.

### Generation of active BCT1 mutants by directed evolution

We also employed a directed evolution approach within a specialized *E. coli* strain lacking CAs (CA-free; Desmarais *et al*., 2019) to evolve functional forms of BCT1. This strain only grows under high levels of CO_2_ or in the presence of a functional bicarbonate transporter or CA (Du *et al*., 2014; Förster *et al*., 2023). By controlling the CO_2_ supply, we determined that a 0.85% (v/v) CO_2_ allowed CA-free to survive for extended periods in liquid culture with slow growth, providing an opportunity for random mutations in the BCT1 plasmid to confer growth advantages. Upon improved growth, cells were transferred to air levels of CO_2_ to increase selection pressure and select functional mutants. The culture was further incubated until it exhibited consistent overnight growth. The duration of the entire process varied from days to weeks. BCT1 plasmids were isolated from single colonies, sequenced and re-transformed into CA-free to confirm the mutations were responsible for the observed growth.

This directed evolution approach led to the generation of two distinct BCT1 mutants (*Figure 4G-H*). In the first, the deletion of the last 450 residues of CmpC (including the regulatory domain) and the first 240 residues of CmpD, resulted in a CmpCD chimera of 263 residues (29 kDa). This mutant also harboured a point mutation in the non-coding intergenic sequence between *cmpA* and *cmpB*. In the second, the deletion of the intergenic space between *cmpC* and *cmpD* produced a CmpCD fusion of 942 residues (105 kDa) that maintained the integrity of both CmpC and CmpD. This mutant also harbored a point mutation in the regulatory domain of CmpC^H409Q^.

### High-throughput screening of BCT1 mutants in CA-free E. coli

A high-throughput complementation plate assay was used to rapidly assess BCT1 function of rational design and directed evolution mutants in CA-free *E. coli*. We screened 72 genetic constructs of BCT1 (*Supplementary Table S2*), of which 14 are shown in *Figure 5*, with each construct identified by a unique identification number. Initially, we confirmed that the unmodified BCT1 construct (GN18) failed to complement CA-free at ambient CO_2_ (0.04%, *Figure 5*). We also evaluated our rational designs, including without a regulatory domain (GN109), phosphorylation mimic (GN113), and half-transporter (GN135). None of these designs supported growth at air (*Figure 5*). However, removing CmpC regulatory domain in our half-transporter design enabled growth at ambient CO_2_ (GN138; *Figure 5*). The addition of small epitope tags to this improved half-transporter design still allowed partial growth at air (GN133; *Figure 5*).

**Figure 5.**
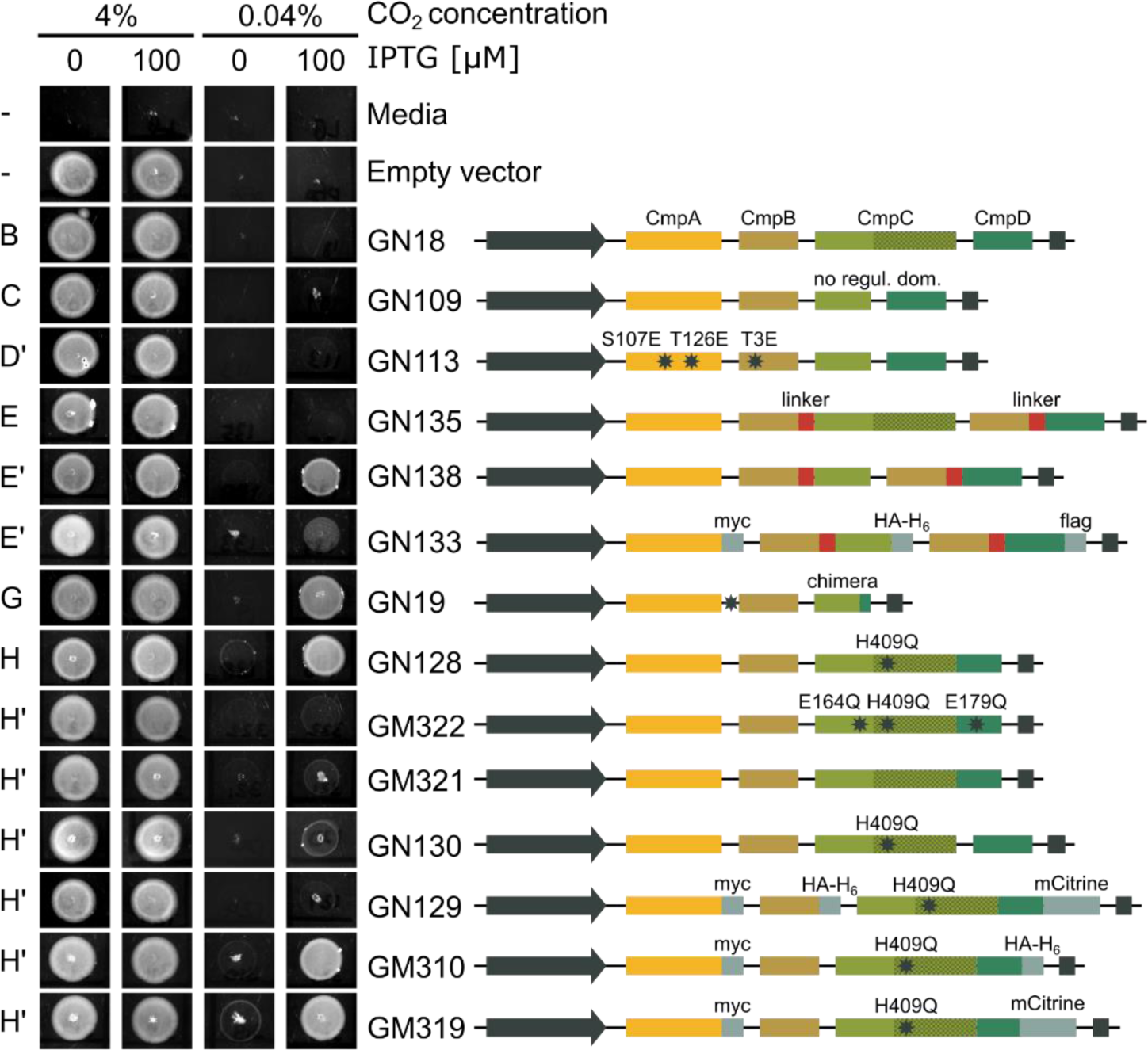
High-throughput spot test screening of BCT1 mutants in CA-free E. coli. Plasmids carrying BCT1 variants, depicted on the right-hand side, were introduced into CA-free *E. coli*. The plasmid backbone used is a Loop-compatible, modified version of pFA31, featuring a LacIQ-pTrc-pLac repressor/promoter cassette (grey arrow) and rrnB T1 & T2 terminator (grey box). On the left-hand side, cultures were plated in 5 µL spots on LB Agar containing 0 or 100 µM IPTG and incubated overnight at 37°C in high (4%) or ambient (0.04%) CO_2_. Successful complementation was achieved when the induced cells (100 µM IPTG) were able to grow at ambient CO_2_ (as observed in the last column). While unmodified BCT1 (GN18) was inactive, seven out of 13 mutants were able to complement CA-free *E. coli* to different extents at ambient levels of CO_2_ (e.g. GN138, GN19, GN128, GM310). The corresponding schematic (see *Figure 4*) to which each plasmid relates to or derives from (indicated by an apostrophe) is presented on the far left as the panel letter from *Figure 4* itsel*f*. The black stars represent point mutations which are labelled, unless falling into a non-coding region (e.g. mutation between *cmpA* and *cmpB* in GN19), to show the change in residues (e.g. H409Q in GN128).

Among the selected BCT1 constructs, two directed evolution mutants (CmpCD chimera [GN19] and CmpCD fusion [GN128]), exhibited successful complementation of CA-free at air. Given its robust complementation ability, we focused our efforts on the CmpCD fusion construct (also containing the H409Q mutation in the regulatory domain of CmpC, *Figure 4H*). Firstly, we demonstrated that the complementation depends on BCT1’s ability to hydrolyse ATP by mutating the catalytic glutamate in CmpC^E164Q^ and CmpD^E179Q^. The ATPase deficient CmpCD fusion failed to complement CA-free (GM322; *Figure 5*). Secondly, we teased apart the influence of the fusion event (between *cmpC* and *cmpD*) and the H409Q mutation in the regulatory domain of CmpC. When the residue Q409 was mutated back into a histidine, the resulting fusion construct failed to complement CA-free (GM321; *Figure 5*). But when a stop codon and an intergenic space were reintroduced between *cmpC^H409Q^* and *cmpD*, the resulting construct weakly complemented CA-free (GN130; *Figure 5*). Finally, we looked at the influence of epitope tags on the CmpCD fusion revealing that while the addition of a tag on CmpA and/or CmpCD fusion had little impact on BCT1 function (GM310, GM319, *Figure 5*; GM315, GM317, *Supplementary Table S2*), a C-terminal tag on CmpB always resulted in a loss of function (GN129, *Figure 5*; GM316, GM318, GM320, *Supplementary Table S2*).

### Functional analysis of selected BCT1 mutants in E. coli

To gain insights into the functional properties of some BCT1 mutants, we conducted H^14^CO_3-_ uptake assays in *E. coli* as described by Förster et al. (2023). Bicarbonate uptake rates were measured for a subset of seven genetic constructs (*Figure 6A*), with the CmpCD fusion exhibiting the highest uptake rate (GN128, 104.2±4.6 nmol.OD_600-1._h^-1^). The addition of a myc tag on CmpA and an mCitrine tag on CmpCD led to a 1.5-fold decrease in uptake rate (GM319, 69.3±10.1 nmol.OD_600-1._h^-1^). Furthermore, replacing mCitrine with HA-H_6_ on CmpCD resulted in a total loss of activity (GM310, 5.5±1.6 nmol.OD_600-1._h^-1^). The improved half-transporter design displayed moderate performance (GN138, 22.5±9.3 nmol.OD_600-1._h^-1^), but the addition of tags reduced the transporter’s activity (GN133, 10±2.3 nmol.OD_600-1._h^-1^) to the same negligible level observed with the unmodified BCT1 (GN18, 10.8±4 nmol.OD_600-1._h^-1^). The CmpCD chimera also exhibited a negligible uptake rate (GN19, 13.4±2.8 nmol.OD_600-1._h^-1^). The kinetic constants were determined for a subset of three constructs which revealed a bicarbonate affinity of approximately 150 µM for both the CmpCD fusion with tags (GM319; K_M_ = 0.17±0.03 mM) and without tags (GN128; K_M_ = 0.12±0.02 mM; *Figure 6B*).

**Figure 6.**
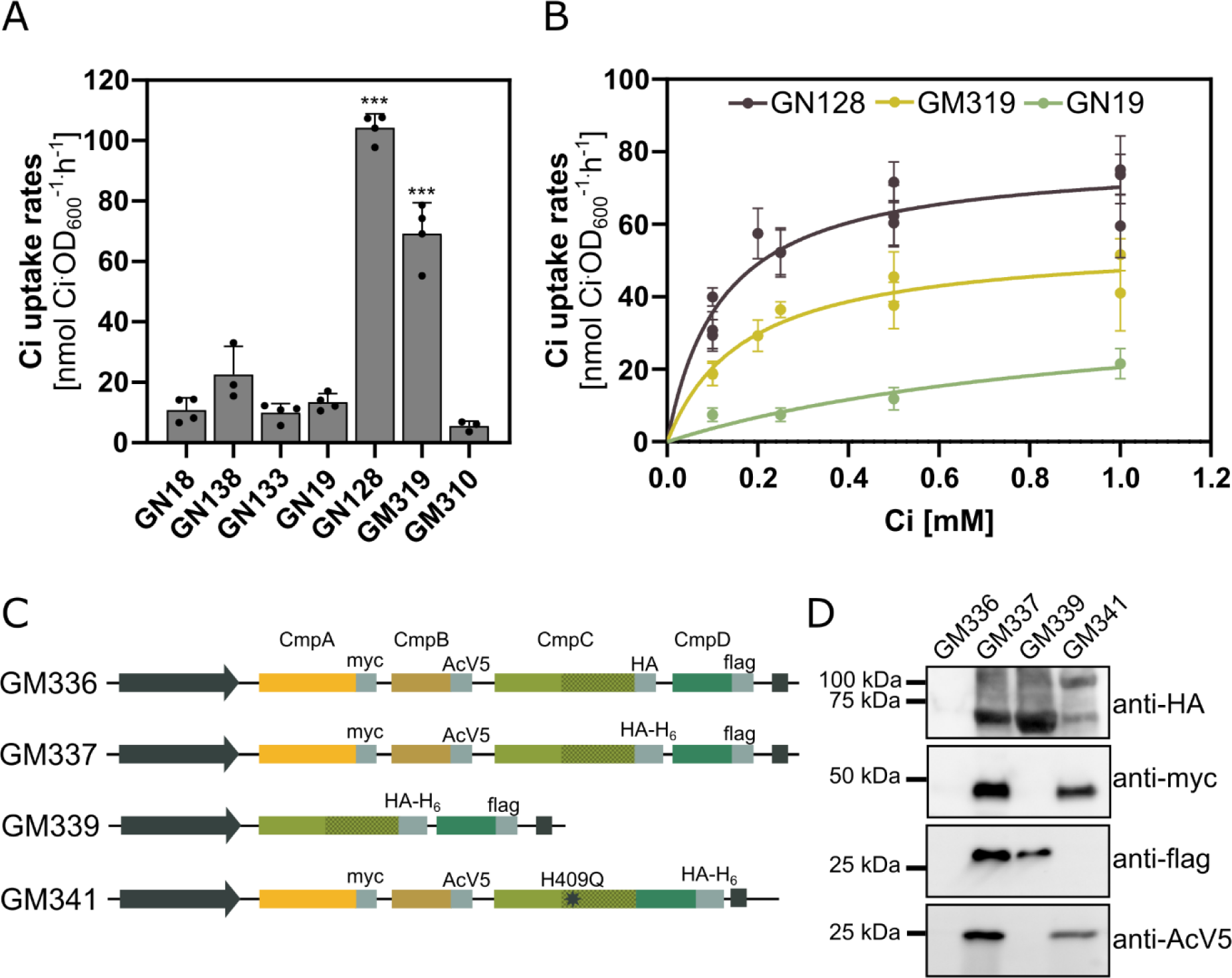
Functional analysis of BCT1 mutants in E. coli by uptake (A-B) and pull-down (C-D) assays. **(A)** Representative bicarbonate uptake rates measured in *E. coli* in presence of 0.5 mM of Ci for a subset of seven BCT1 mutants. The constructs used here are depicted in *Figure 5.* The values obtained with an empty vector, representing background CO_2_ diffusion, have been subtracted. Statistical differences across mutants were assessed with a one-way ANOVA followed by pairwise multiple comparisons. Asterisks are an indication of the *P*-value (\*\*\**P* < 0.001) relative to the unmodified BCT1 (GN18). Mean ±SD (n=4). **(B)** Representative bicarbonate uptake curves for selected BCT1 mutants measured in *E. coli*. The Michaelis-Menten equation was fitted to the data by non-linear regression to obtain the maximal velocity (V_MAX_) and affinity constant (K_M_). Individual data points represent the mean of 4 technical replicates at each bicarbonate concentration (±SD). **(C)** Depiction of the constructs used for IMAC pull-downs. **(D)** Western blot of the IMAC eluate showing co-purification of the BCT1 complex in *E. coli*. Loaded 10 µL of the concentrated eluate. Note that GM341 lacks a flag tag because CmpD is fused to CmpC and is detected with HA-H_6_ around 107 kDa.

We also explored the assembly of the BCT1 complex in *E. coli*. To facilitate this assessment, each BCT1 protein was tagged with a small epitope (*Figure 6C*). CmpC, the bait protein, was purified by virtue of its C-terminal hexa-histidine tag using Immobilized Metal Affinity Chromatography (IMAC), with the expectation that interacting proteins (prey) would co-purify. A negative control involved using a BCT1 construct with identical tags, except for the absence of the hexa-histidine tag on CmpC (GM336). This control confirmed the effectiveness of the column washes, as no signal was detected in the eluate fraction for GM336 (*Figure 6D*). The IMAC pull-down was then repeated with three different BCT1 constructs. The eluate of the unmodified BCT1 (GM337) contained all four proteins, indicating that the presence of a tag on CmpB does not obstruct transporter assembly. The NBD-only construct (GM339) revealed that CmpC and CmpD can directly interact without necessitating CmpB to form a heterodimer. Lastly, in GM341, where CmpC is fused with CmpD, the interaction with CmpB persisted, with both CmpB and CmpA detected in the eluate fraction.

### Functional analysis of selected BCT1 mutants in Arabidopsis

Six BCT1 genetic constructs were adapted for plant expression and introduced into the Arabidopsis *βca5* mutant (*Supplementary Figure S5*). *βca5* lacks the plastidial carbonic anhydrase βCA5, and like CA-free *E.coli*, is unable to grow at air unless expressing a functional bicarbonate transporter or a plastid-localized CA (Weerasooriya *et al*., 2022; Förster *et al*., 2023). However, when transformed into *βca5,* none of the tested BCT1 constructs, including unmodified (GN23), no regulatory domain (GN24), phosphorylation mimic (GN55), half-transporter (GN64), half-transporter with CmpC_263_ (GN65), and CmpCD fusion (GN139), restored *βca5* growth at air (*Figure 7*).

**Figure 7.**
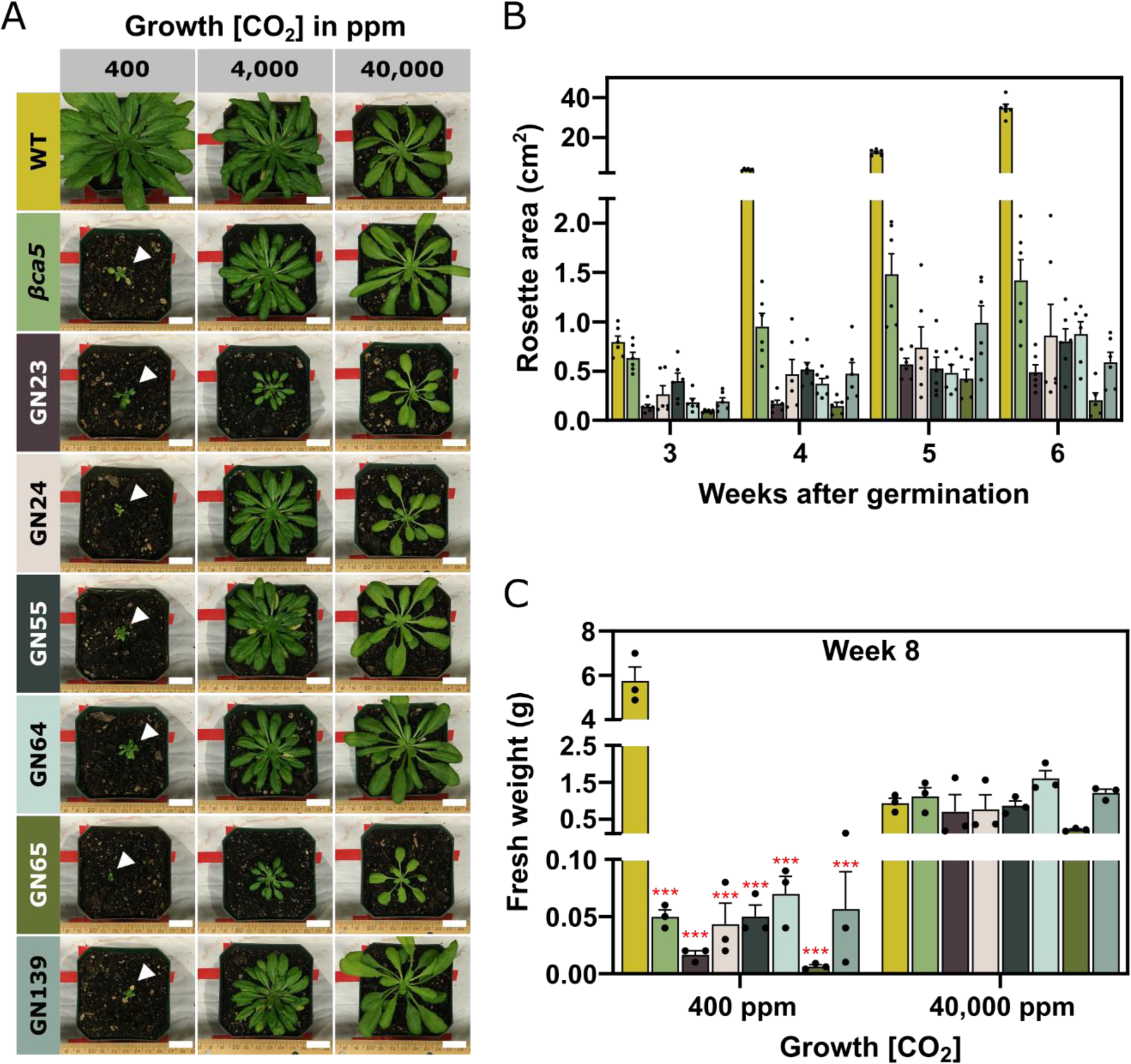
Complementation of the Arabidopsis βca5 mutant. Plants were grown at ambient (400 ppm), high (4,000 ppm) or very high (40,000 ppm) CO_2_ concentrations to assess the complementation ability of various BCT1 mutants. The genetic constructs used to transform the *βca5* mutant are depicted in *Supplementary Figure S5.* Colours are used consistently between the three panels. **(A)** Images of wild-type (WT; Col-0) and transformed *βca5* mutant (SALK_121932) *A. thaliana* plants eight weeks after germination. The images are representative of six plants. Scale bar shown is 2 cm long. **(B)** Overhead images of plants grown at ambient CO_2_ were taken weekly, and rosette areas were measured using the PhenoImage and ImageJ software. Mean ±SE (n=6). **(C)** Plants were harvested for fresh weight eight weeks after germination. Statistical differences across genotypes were assessed with a one-way ANOVA followed by pairwise multiple comparisons between plants at each CO_2_ concentration. Red asterisks are an indication of the *P*-value relative to WT (\**P* < 0.05; \*\**P* < 0.01; \*\*\**P* < 0.001). Mean ±SE (n=3).

To further our analysis, the two half-transporter constructs (GN64, GN65), and CmpCD fusion (GN139) were introduced into wild-type (WT) Arabidopsis. These plants were then grown on air levels of CO_2_ (400 ppm) or low CO_2_ (200 ppm) to determine whether the BCT1 constructs might enhance growth. None of these constructs enhanced growth of WT Arabidopsis when grown at these CO_2_ levels (*Figure 8*, *Supplementary Figure S6*). In addition, the mature leaves of the transformed plants displayed lower or similar CO_2_ assimilation rates (*A/Ci* curves; CO_2_ assimilation rate as a function of intercellular CO_2_) as compared to WT, with CO_2_ compensation points unchanged or higher than WT (*Supplementary Table S3*).

**Figure 8.**
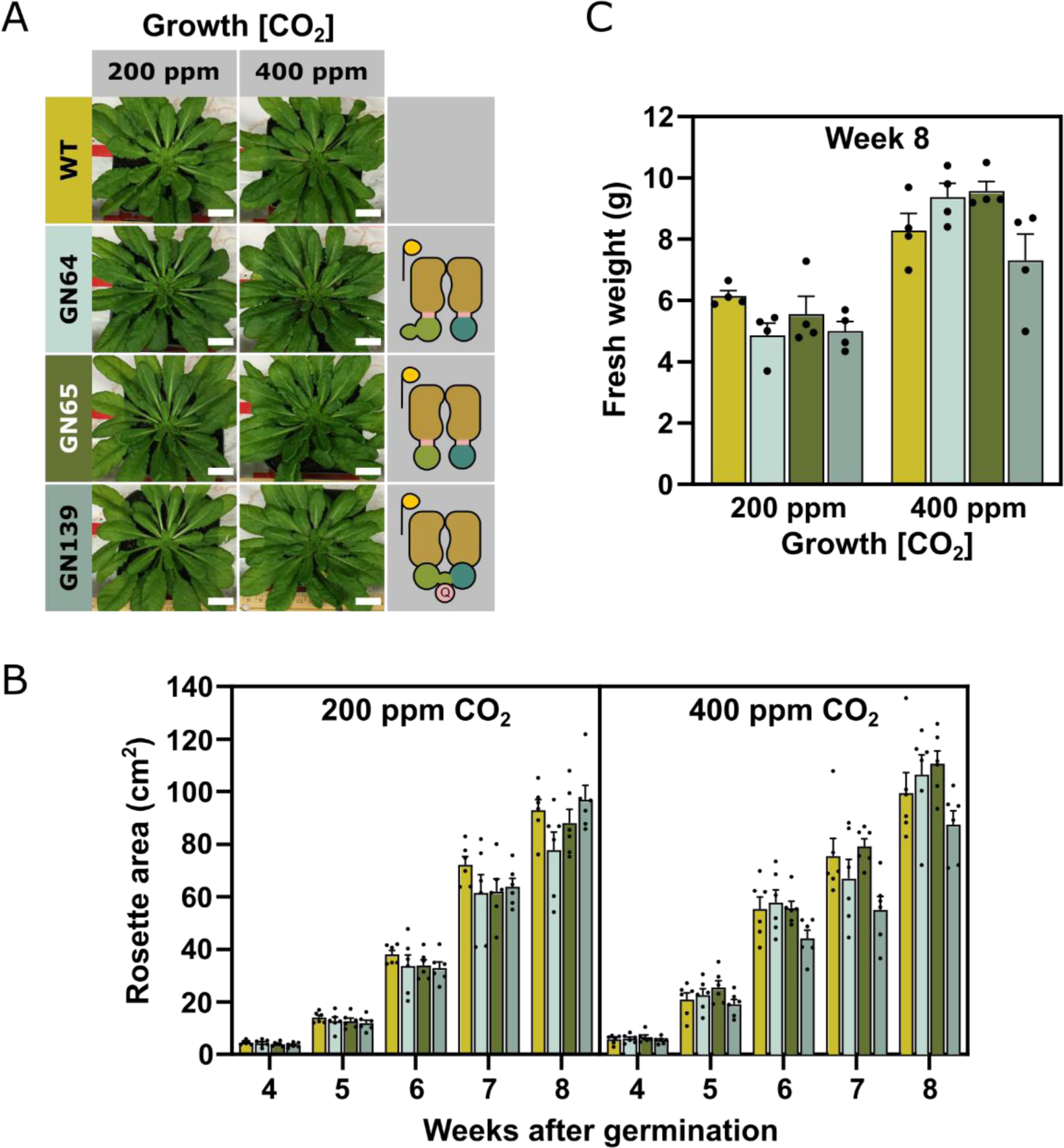
Functional analysis of BCT1 transformants in WT Arabidopsis. (**A**) Images of 8-week-old *A. thaliana* (Col-0) plants transformed with three BCT1 constructs (GN64, GN65, GN139). The plants were grown at ambient (400 ppm) or reduced CO_2_ concentrations (200 ppm). The images are representative of six plants. Scale bar shown is 2 cm long. Depictions of BCT1 mutants is on the right-hand side. GN64 and GN65 are translational fusions of CmpBC and CmpBD (reflecting a half-transporter design, Ford *et al*., 2019) and GN139 is a CmpCD fusion obtained by directed evolution. In the half-transporter design, GN64 harbors full-length CmpC while in GN65 CmpC has no regulatory domain (i.e., CmpC_263_). BCT1 subunit colours are as described in *Figure 4A*.(**B**) Overhead images of the plants were taken weekly, and rosette areas were measured using the PhenoImage and ImageJ software. Mean ±SE (n=6). **(C)** Plants were harvested for fresh weight 8 weeks after germination. Statistical differences across genotypes were assessed with a one-way ANOVA followed by pairwise multiple comparisons between plants at each CO_2_ concentration. No statistical difference was recorded. Mean ±SE (n=4). Colours are used consistently between the three panels and are the same as used in *Figure 7*.

## Discussion

In this study, we present compelling evidence supporting the independent functional evolution, and precise subcellular targeting of a complex cyanobacterial bicarbonate transporter. Our primary objective was to introduce a functional Ci transporter into plants, aiming to enhance CO_2_ assimilation in C_3_ crops (Price *et al*., 2013). Previous research in this field predominantly focused on simpler single or dual-gene bicarbonate pump systems, often encountering difficulties related to targeting or additional ion requirements for function (Pengelly *et al*., 2014; Atkinson *et al*., 2016; Rolland *et al*., 2016; Uehara *et al*., 2016, 2020; Nölke *et al*., 2019; Rottet *et al*., 2021; Förster *et al*., 2023). The successful integration of the *Chlamydomonas* passive channel LCIA into C_3_ plant chloroplasts was previously accomplished; however, this transporter’s inherent characteristics as a passive channel limit its capacity for high-rate bicarbonate transport (Atkinson *et al*., 2016; Nölke *et al*., 2019; Förster *et al*., 2023).

For the first time, we addressed dual challenges described in previous reports: independently achieving transporter functionality, and correct subcellular localization of a foreign bicarbonate transporter in plants. Notably, we directed the ABC transporter BCT1 to the chloroplast envelope, a complex task given its four subunits, each needing precise localization (*Figure 1*). BCT1 was previously reported to be inactive in *E. coli*, potentially due to unknown regulatory mechanisms likely present in its native cyanobacterial cellular environment (Du *et al*., 2014). To remove regulatory requirements, BCT1 was engineered, and its functionality assessed in a specialized *E. coli* strain. Despite these complexities, BCT1 possesses favourable attributes, including a high affinity for bicarbonate and the reliance on ATP as its sole power source (Omata *et al*., 1999*b*,*a*), eliminating the need for co-transported ions, as is the case for the single gene transporters SbtA and BicA (Price *et al*., 2004, 2008).

The ability to import nuclear-encoded proteins into chloroplasts has a broad application to the majority of globally important crops. The assembly of a multi protein membrane complex in a heterologous system is a significant engineering challenge. It requires the components to be co-localized and for the membrane proteins to be inserted in the correct orientation (Wojcik and Kriechbaumer, 2021). Factors such as stoichiometry and chaperones may also have to be considered (Barrera *et al*., 2009; Bae *et al*., 2013; Hallworth *et al*., 2013; Thornell and Bevensee, 2015). We found that the BCT1 complex assembled in *E. coli* (*Figure 6*) and in *N. benthamiana* (*Figure 3*). A critical observation was the recruitment of CmpC and CmpD to the chloroplast IEM when co-expressed with CmpB, which suggests that CmpB is oriented correctly in the membrane irrespective of which leader sequences was used (*Figure 3* and *Supplementary Figure S3*). This is not only essential for the complex formation but also guarantees the intended direction of transport. Notably, we observed that not all leader sequences were equally effective at targeting BCT1 component proteins to the correct locations within the chloroplast. For example, *At*MGD1 failed to target CmpA to the IMS (*Supplementary Figure S1*). Additionally, while the *At*RecA leader sequence proved highly efficient for directing CmpD to the stroma, it could not efficiently deliver CmpC, possibly due to steric hindrance issues (Figure 2; Köhler *et al*., 1997; Shen *et al*., 2017). As more leader sequences become available, our toolkit for subcellular targeting will expand, and the use of modular cloning will enable rapid screening of additional sequences.

Initially, native BCT1 was inactive in *E. coli* (Figures 5 and 6; Du *et al*., 2014). We hypothesized the lack of function was due to the absence of regulatory factors in heterologous systems (e.g. specific activation kinases; Spät *et al*., 2021). To overcome this problem, we used two approaches. Logical changes were made to the proteins by rational design, and directed evolution was employed to evolve active forms of BCT1 (*Figures 4* and *5*). Directed evolution led to large changes such as the fusion of the two NBDs in a CmpCD fusion. With rational design, we explored the fusion of the TMD with each NBDs in the CmpBC and CmpBD half-transporter design (Theodoulou and Kerr, 2015; Ford *et al*., 2019). In both approaches we obtained some level of activity, suggesting that subunit stoichiometry plays an important role for the functionality of BCT1, as protein fusion likely altered the CmpB:CmpC/D ratio.

We also hypothesised that eliminating the CmpC regulatory domain could produce an active transporter. While this rationally designed form, CmpC_263_, did not show the predicted activity, directed evolution produced a CmpCD chimera which had measurable activity in the absence of this regulatory domain (*Figures 4*). Additionally, a CmpCD fusion, which was the best-performing mutant, harboured a point mutation in the regulatory domain of CmpC^H409Q^. This mutation played a more significant role than the fusion event itself. However, both the mutation and fusion were found to be necessary for achieving maximal activity of the transporter. A multiple sequence alignment (*Supplementary Figure S7*) showed that residue H409 in CmpC corresponds to putative ligand-binding residues in NrtA (H196) and CmpA (Q198; Koropatkin *et al*., 2006, 2007). Considering this, we speculate that the H409Q mutation might interfere with ligand binding in some manner. Further research is needed to understand the role of H409 but with eight potential binding sites identified in CmpA and NrtA, we predict there are still many unexplored rational designs that could lead to an improved functionality of BCT1.

Based on functional modification of BCT1 through rational design and directed evolution, and the ability to successfully target BCT1 components to their destinations within the chloroplasts, we generated transgenic Arabidopsis lines expressing several modified BCT1 constructs (*Supplementary Figure S5*). Notably, none of these, either in *βca5* mutant (*Figure 7*) or in WT plants (*Figure 8*), displayed phenotypes consistent with bicarbonate uptake into the chloroplast. Also, the expected decrease in CO_2_ compensation point was not apparent (*Supplementary Table S3*). Bicarbonate uptake into the chloroplast should enhance chloroplastic CO_2_ concentrations, elevating Rubisco carboxylation even at low ambient CO_2_ supply (Price *et al*., 2011). The lack of a CO_2_ compensation point reduction in our BCT1 lines and the failure of these constructs to enhance the growth of plants indicates that BCT1 is not significantly changing chloroplast Ci uptake in these plants.

We hypothesise that further evolution and refinement of function of BCT1 in the CA-free *E. coli* system may be required to deliver improved function *in planta*. Notably, the large sequence changes observed using directed evolution in this study, and the similarity between evolved outcomes and some of the rational designs, highlights two things. Firstly, that well-considered rational design approaches using known variation in evolution of ABC transporter systems (e.g., half-transporter protein fusion arrangement) is a valid approach to modify this type of transporter. Secondly, our directed evolution approach enabled the generation of large and unexpected changes in sequence length and gene fusion that would have not been found in the screen of a sequence variant library. We are therefore encouraged that a combination of rational design and directed evolution of both existing chloroplast membrane proteins and bacterial bicarbonate uptake systems will allow significant progress in enabling the elevation of chloroplastic Ci using synthetic biology tools. In combination with high throughput DNA assembly technologies and plant-based platforms that enable functional testing, we expect significant progress toward this goal.

## Supplementary data

The following supplementary data are available online.

*Figure S1* Targeting of CmpA to the chloroplast intermembrane space.

*Figure S2* Targeting of CmpB to the chloroplast inner envelope membrane.

*Figure S3* Orientation of CmpB in the inner envelope membrane.

*Figure S4* Structure of CmpC.

*Figure S5* Genetic constructs screened in Arabidopsis *βca5* mutant.

*Figure S6* Rosette area and assimilation rate in transgenic Arabidopsis.

*Figure S7* Sequence alignment and corresponding WebLogo conservation sequence of CmpC, CmpA, NrtA and NrtC from β- and α-cyanobacteria.

*Table S1* List of primers used in this study.

*Table S2* List of constructs used in this study.

*Table S3* CO_2_ compensation points for BCT1 transformants in WT Arabidopsis.

## Materials and methods

### Construction of BCT1 expression vectors

DNA plasmid constructs were produced using type IIS cloning strategies adapted from Golden Gate cloning and Loop Assembly (Engler *et al*., 2014; Pollak *et al*., 2019). BCT1 genes were amplified from *Synechococcus sp*. PCC7942 and domesticated to remove type IIS restriction sites. Primers were designed around the gene of interest with *Bpi*I recognition sites and an appropriate 4-bp overhang (*Supplementary Table S1*). Polymerase chain reaction (PCR) was performed using Phusion^TM^ High-Fidelity DNA Polymerase (ThermoFisher Scientifc, USA), the bands of desired sizes were gel-purified using Promega Wizard® SV Gel and PCR Clean-Up System (Promega, USA). PCR fragments were assembled into the Universal Level 0 vector (pAGM9121) under cyclical digestion and ligation condition (37°C for 3 minutes, 16°C for 4 minutes for 25 cycles) followed by heat inactivation (50°C for 5 minutes, 80°C for 5 minutes). The same cyclical digestion and ligation condition with heat inactivation was performed when assembling Level 1, Level 2 and Level 3 constructs, but with different restriction enzymes and acceptor plasmids. While BbsI-HF® (New England BioLabs, USA) was used for Level 0 assembly, BsaI-HF®v2 (New England BioLabs, USA) was used for Level 1 and 3 assembly and SapI (New England BioLabs, USA) for Level 2 assembly. Acceptors were pOdd1-4 (pCk1-4, Addgene plasmids # 136695-136698) for Level 1 and 3 and pEven1-4 (pCsA-E, Addgene plasmids # 136067-136070) for Level 2. To optimize BCT1 expression in *E. coli*, the low copy number pFA31 backbone (Addgene plasmid #162708; Flamholz *et al*., 2020) was modified into two terminal acceptors compatible with Loop Assembly (pFA-Odd and pFA-Even). For the ‘half-transporter’ designs, flexible linkers were adapted from BBa_K365005, BBa_K157013, BBa_K157013, BBa_K157009 (iGem Standard Biological Parts, http://parts.igem.org/). QIAprep Spin Miniprep Kit (Qiagen, USA) was used for all plasmid purification and construct sequences were confirmed by Sanger sequencing (Macrogen Inc., Seoul South Korea). Primers used for assembling and checking the different constructs can be found in *Supplementary Table S1*.

### Plant growth conditions

*N. benthamiana* plants used for infiltration were grown under 400 μmol photons m^-2^ s^-1^ light intensity, 60% relative humidity, a 16 h light/8 h dark photoperiod and 25°C day/20°C night temperatures. Only the 1^st^, 2^nd^ and 3^rd^ true leaves from 4 to 5-week-old plants were kept for infiltration, while the rest were discarded. The plants were germinated and grown on pasteurized seed raising mix supplemented with 3 g/L Osmocote Exact Mini.

WT (Col-0) and *βca5* mutant (SALK_121932; obtained from TAIR) *A. thaliana* plants were used for transformation experiments with the various BCT1 constructs. The plants were grown in Metro-Mix 830 (Sun Gro Horticulture, Agawam, MA, USA) with 100 μmol photons m^-2^ s^-1^ light intensity under short days (8 h light/16 h dark). WT plants were grown in ambient (400 µL L^−1^ CO_2_) and reduced CO_2_ (200 µL L^−1^ CO_2_) conditions, while *βca5* mutants were supplemented with high CO_2_ (0.4% v/v CO_2_ or 4000 µL L^−1^ CO_2_) or very high CO_2_ (4% v/v CO_2_ or 40000 µL L^−1^ CO_2_) to allow normal growth. The plants were maintained with distilled H_2_O and a 1:3 dilution of Hoagland’s nutrient solution (Epstein and Bloom, 2005).

### Agroinfiltration of Nicotiana benthamiana leaves

Constructs for BCT1 localization studies (*Supplementary Table S2*) were transiently expressed in 4-5 week old *N. benthamiana* leaf tissue via Agrobacterium infiltration, as described previously (Rolland, 2018). Briefly, *A. tumefaciens* GV3101 (pMP90) were transformed with BCT1 constructs and grown in lysogeny broth (LB) media supplemented with 25 µg mL^−1^ rifampicin, 50 µg mL^−1^ gentamycin and 50 µg mL^−1^ kanamycin or 100 µg mL^−1^ spectinomycin for 24 hours at 28°C and 200 rpm. A vector encoding the tomato bushy stunt virus P19 protein was used to inhibit post-transcriptional gene silencing and to enable the expression of our constructs of interest (Roth *et al*., 2004). For each infiltration, p19 culture was mixed with each construct of interest at an OD_600_ of 0.3 and 0.5, respectively. A p19-only control was prepared to an OD_600_ of 0.8 as a negative control. All cells for infiltration were pelleted at 2,150 g for 8 minutes and resuspended in 5 mL of infiltration solution (10 mM MES pH 5.6, 10 mM MgCl_2_, 150 µM acetosyringone). The solutions were incubated at room temperature for 2 hours with occasional swirling, then infiltrated into the abaxial surface of 4-week-old *N. benthamiana* leaves using a 1 mL slip tip syringe. Infiltrated plants were grown for another 3 days before protein expression was assessed via confocal microscopy.

### Agrobacterium-mediated transformation of Arabidopsis thaliana

BCT1 plant expression vectors were transformed into *Agrobacterium tumefaciens* strain GV3101. Cultures were grown in LB media with antibiotics (30 µg mL^−1^ gentamycin, 10 µg mL^−1^ rifampicin, and 50 µg mL^−1^ kanamycin). *Arabidopsis* plants were transformed using the method described by (Zhang *et al*., 2006). A 5 mL starter culture of *A. tumefaciens* was grown in LB media with antibiotics overnight at 28°C. This starter culture was used the following morning to propagate a larger 250 mL *A. tumefaciens* culture overnight at 28°C. The next day, the cells were harvested by centrifugation at 4,000 g for 10 minutes. The pelleted cells were resuspended in freshly prepared 5% (w/v) sucrose solution with 0.02% (v/v) of Silwet L-77. The resuspended cultures were generously applied to the *Arabidopsis* flower buds using transfer pipettes. Afterwards, the plants were placed sideways into the trays and were covered and allowed to recover in darkness overnight. Following recovery, the plants were grown in 21°C in continuous light. Mature seeds were collected from plants and positive transformants were selected on soil by spraying seedlings with a 1:2000 dilution of BASTA (AgrEvo, Berlin, Germany). The presence of the transgene was also confirmed via gene-specific PCR for *cmpA* with the primer pair CmpA-F1 and CmpA-R1 and for the *bar* gene with the primer pair Basta-F and Basta-R (*Supplementary Table S1*). DNA was extracted using the protocol described by (Edwards *et al*., 1991). Namely, about 20 mg of plant tissue was ground using micropestles in 1.5 mL centrifuge tubes. These were further macerated in 400 µL of extraction buffer (200 mM Tris-HCl pH 7.5, 250 mM NaCl, 25 mM EDTA, 0.5% [w/v] SDS). The samples were then centrifuged at 13,000 g for 5 minutes, and the supernatant was collected into a new tube. An equal volume (∼400 µL) of isopropanol was added and mixed to the supernatant. The resulting mixture was again centrifuged at 13,000 g for 5 minutes. The resulting supernatant was discarded afterwards, and the pellet was allowed to air dry. After drying, the pellet was dissolved in 50 µL of 1X TE buffer (10 mM Tris-HCl, 1mM Na_2_EDTA, pH 8.0) and was used for subsequent confirmation of transformation.

### Confocal microscopy

Confocal laser microscopy was performed on *N. benthamiana* infiltrated with BCT1 constructs at 3-4 dpi (days post-infiltration). In 3 independent experiments, leaf disks or protoplasts (Rolland, 2018) were observed and several images taken, using a Leica SP8 confocal laser microscope, a 63x water immersion objective (NA= 1.2), PMT detectors and the Leica Application Suite X software package. Confocal microscope settings for the detection of chlorophyll (λ_ex_=488 nm or 514 nm, λ_em_=650-690 nm), mCitrine (λ_ex_=514 nm, λ_em_=520-540 nm), and mNeon (λ_ex_=488 nm, λ_em_=512-530 nm) were as described previously (Stoddard and Rolland, 2019).

### Bacterial strains and growth conditions

*E. coli* CA-free strain, kindly provided by Dave Savage (Desmarais *et al*., 2019), was used for directed evolution and complementation assay. *E. coli* DH5α strain was used for cloning and protein expression for IMAC (immobilized metal affinity chromatography) purification. Unless otherwise stated, bacteria were grown at 37°C in LB media (10 g/L tryptone, 10 g/L NaCl and 5 g/L yeast extract), supplemented with 15 g/L agar for solid media on plates. For culturing transformants with spectinomycin, ampicillin and kanamycin resistant genes, media were supplemented with 100 µg mL^−1^, 100 µg mL^−1^ and 50 µg mL^−1^ of the antibiotics respectively.

### Directed evolution of BCT1 in CA-free E. coli

By controlling the CO_2_ supply, we determined that a 0.85% (v/v) CO_2_ allowed CA-free to survive for extended periods in liquid culture, providing an opportunity for random mutations in the BCT1 plasmid to confer growth advantages.

A starter culture of CA-free harboring unmodified BCT1 genes in pEven1 backbone (GM186) was prepared by growing the cells from a glycerol stock in LB medium supplemented with 100 µg mL^−1^ spectinomycin at 37°C in the presence of 4% CO_2_ for approximately 18 hours. This starter culture was then diluted 100 µL into 5 mL of liquid media consisting of M9 minimal medium supplemented with 1% LB, 100 µg mL^−1^ spectinomycin, and 20 µM isopropyl β-D-1-thiogalactopyranoside (IPTG). The cultures were incubated at 37°C with agitation at 120 rpm under a 0.85% CO_2_ atmosphere. Regular subculturing in fresh media were performed while maintaining permissive CO_2_ conditions, until the cultures were able to fully grow overnight.

The overnight culture was diluted 100-fold and placed under ambient CO_2_ conditions (0.04%). Regular subculturing was again performed until the cultures reached a dense overnight growth. From the cultures that grew under ambient CO_2_, 100 µL was plated on solid LB media supplemented with 100 µg mL^−1^ spectinomycin and 100 µM IPTG, and incubated at ambient CO_2_ for 18 hours. Eight single colonies were selected and cultured, and their plasmid DNA was extracted using the QIAprep Spin Miniprep Kit (Qiagen, USA). The pDNA from these colonies was pooled together and used to retransform new CA-free cells. The transformed cells were then plated onto LB agar supplemented with 100 µg mL^−1^ spectinomycin and 100 µM IPTG and incubated at ambient CO_2_ for 18 hours. Growth confirmed that the mutation(s) conferring the advantage at air was carried by the plasmid and not in the genome of the CA-free strain. Twelve colonies were selected from this plate, and the pDNA was isolated from each colony for DNA sequencing (Macrogen Inc., Seoul, South Korea).

For the isolation of the strain harbouring the CmpCD chimera, cells were subcultured four times at 0.85% CO_2_ and three times at air before it grew overnight. For the isolation of the strain harbouring the CmpCD fusion, cells were subcultured six times at 0.85% CO_2_ before it grew overnight at air.

### Complementation assay in CA-free E. coli

We developed a high-throughput complementation assay to rapidly assess BCT1 function in CA-free. This involved cultivating liquid cultures at 4% CO_2_ for 6 hours at 37°C, spotting 5 µL onto four plates of LB agar with or without 0.1 mM IPTG, and incubating them overnight in selective (air, 0.04% CO_2_) or permissive conditions (4% CO_2_). To mitigate the negative growth effects caused by BCT1 overexpression, a modified plasmid backbone with lower copy number was employed in this assay (pFA-Odd and pFA-Even). After the overnight growth, the plates were imaged with a Bio-Rad ChemiDoc XRS+ imaging system (Bio-Rad, USA) under white epifluorescence.

### Bicarbonate uptake assay in E. coli

Inorganic carbon uptake assays were carried out as described by Förster et al. (2023) with some modifications. The assays were performed in CA-free, and the cultures were induced with 100 µM IPTG. The assay buffer used consisted of 20 mM bis-tris propane-H_2_SO_4_ supplemented with 0.5 mM glucose and 1 µM CaCl_2_ with a pH of 7.5.

To prepare the cells for the assay, they were first washed twice with the assay buffer. Subsequently, the cells were incubated for ten minutes before undergoing a third round of washing with the assay buffer. Following the washing steps, the assay was performed according to the protocol outlined in Förster et al. (2023). This method allowed us to measure the rates of bicarbonate uptake and determine kinetic parameters such as the Michaelis constant (K_M_) and maximum velocity (V_MAX_) for bicarbonate transport.

### Protein induction and Immobilised Metal Affinity Chromatography purification

Overnight cultures from glycerol stocks were used to inoculate 40 mL of LB medium supplemented with 100 µg mL^−1^ spectinomycin to an optical density (OD_600_) of 0.1-0.2. Cultures were grown at 37°C until OD_600_ reached 0.4-0.6. To induce protein expression, IPTG was added to a final concentration of 50 µM. Cultures were returned to grow at either 37°C for two to three hours or 28°C for four to five hours. To prepare the cells for IMAC purification, the OD_600_ of each culture was measured and used to normalize the number of cells to pellet. The cell pellets were then harvested by centrifugation at 4,800 g for 10 minutes at 4°C and subsequently stored at -20°C until further use.

Cell pellets were resuspended with 1 mL of lysis buffer [5% (v/v) glycerol, 50 mM HEPES pH 8.0, 50 mM NaCl, 1% (v/v) bacterial protease inhibitor cocktail (P8849, Sigma, USA) added fresh before using] and incubated with 1 µL of rLysozyme solution (71110-6000KU, EMD Millipore Corp, USA) for 30 minutes on ice. The suspension was topped up to 5 mL with lysis buffer supplemented with 12.5 mM CaCl_2_, 25 mM NaHCO_3_, 6.25 mM MgCl_2_, 6.25 mM ATP, 6.25 mM Na_3_VO_4_ and 5mM imidazole before lysing with three passes through the Emulsiflex (Avestin, USA) at 60 psi. Lysates were incubated with 1% (w/v) n-Dodecyl-β-D-Maltoside (DDM) detergent with constant gentle rotating at 4°C for 30 minutes, and then clarified via passing through Millex® -GP Fast Flow & Low Binding Millipore Express® PES Membrane 0.22 µm syringe filter unit (SLGP033RS, Merck Millipore, USA).

Each clarified lysate was incubated with 500 µL bed volume of Profinity^TM^ IMAC Ni-Charged Resin (156-0135, Bio-Rad, USA) in Poly-Prep® Chromatography Columns (731-1550, Bio-Rad, USA), pre-washed with 7 mL of binding buffer [5% (v/v) glycerol, 50 mM HEPES pH 8.0, 50 mM NaCl, 10 mM imidazole], under gentle inversion at 4°C for 1 hour. The resin with bound proteins was washed four times with 2.5 mL wash buffer [5% (v/v) glycerol, 50 mM HEPES pH 8.0, 50 mM NaCl, 1% (w/v) DDM, 10 mM CaCl_2_, 20 mM NaHCO_3_, 5 mM MgCl_2_, 5 mM ATP, 5 mM Na_3_VO_4_, 20 mM imidazole] by gravity flow. Proteins were eluted with 2 mL elution buffer [5% (v/v) glycerol, 50 mM HEPES pH 8.0, 300 mM NaCl, 1% (w/v) DDM, 250 mM imidazole]. Prior to SDS-PAGE analysis, eluates were concentrated ∼ 20 times by trichloroacetic acid (TCA) precipitation via addition of 200 µL of 0.15% (w/v) sodium deoxycholate and 200 µL of 72% (w/v) TCA solution. The mixtures were vortexed and incubated at ambient temperature for 5 minutes before being pelleted at 20,238 g for 8 minutes. Protein pellets were resuspended in 120 µL of resuspension buffer (Laemmli sample buffer, 50 mM DTT, 3.84 % (w/v) SDS, 400 mM Tris pH 7.4, 150 mM NaOH, pH 10), kept at 4°C for overnight. For longer storage, samples were kept at -20°C.

### SDS-PAGE and Western blotting

Protein samples were mixed with gel loading buffer (Laemmli Sample Buffer, 50 mM DTT), boiled at 95°C for 10 minutes, centrifuged for 2 minutes before running through 4-20% mini Protean TGX Stain Free Gel (BioRad, USA). The separated proteins were transferred to Immobilon®-P PVDF membrane (Merck Millipore, USA) and probed with primary antibodies overnight. Antibodies used for probing membrane include polyclonal anti-GFP antibody produced in rabbit (1:2000 dilution, abcam, USA), monoclonal anti-FLAG M2 antibody produced in mouse (1:2000 dilution, Sigma-Aldrich, USA), monoclonal anti-c-Myc antibody produced in mouse (1:5000 dilution, Sigma-Aldrich, USA), monoclonal anti-HA antibody produced in mouse (1:5000 dilution, Sigma-Aldrich, USA) and monoclonal anti-AcV5 tag antibody produced in mouse (1:5000 dilution, abcam, USA). Probed membrane was washed with TBS-T thrice before incubating with alkaline phosphatase-conjugated anti-mouse secondary antibody (1:10000 dilution, Sigma, USA) or goat anti-rabbit IgG (H+L) secondary antibody (1:5000 dilution, Invitrogen, USA) for one hour. The blot was washed as before and then visualised with Attophos Substrate Kit (Promega, USA) using the Bio-Rad ChemiDoc XRS+ system.

### Physiological measurements on Arabidopsis

Images of different Arabidopsis genotypes were taken weekly, and rosette areas were measured as pixel area using the PhenoImage software (Zhu *et al*., 2021) and Fiji ImageJ (Schindelin *et al*., 2012). Rosette areas were measured on six plants per line. Four plants were later harvested for measuring fresh weights. Measurements of photosynthetic parameters were conducted using a LI-COR LI-6800 system (Lincoln, NE). Plants grown for rosette area and biomass measurements were also used for photosynthesis measurements. CO_2_ response curves for assimilation (A; µmol m⁻² s⁻¹) in response to intercellular CO_2_ (Ci; µmol mol^-1^) curves were generated from 50 to 1700 µmol mol^-1^ ambient CO_2_.

### Data visualization and statistical analysis

Growth and physiological parameter data were initially visualized using the R software environment and the ggplot2 package (Wickham, 2016; R Core Team, 2019). One-way ANOVA followed by Tukey’s multiple comparisons test was performed using GraphPad Prism version 10.0.3 for Windows, GraphPad Software (Boston, Massachusetts USA, www.graphpad.com).

## Acknowledgements

The authors thank Hanjun Sun for early protocol development for the directed evolution system in CA-free. We also thank Sacha B. Pulsford and Wei Yih Hee for contributing to the design and construction of DNA parts. OpenAI’s ChatGPT 3.5 was used for prose editing.

## Author contributions

GDP, JVM, BML, and SR: conceptualization; GDP, and JVM: project administration; SR, LMR, SY, SYP and ICP: methodology; SR, LMR, ICP, XW, HSM and NDN: formal analysis; SR, LMR, ICP, SYP, SY, XW, HSM and HNW: investigation; BML, SR, JVM and GDP: supervision; SR, NDN and BML: visualization; SR, LMR, SYP, SY, ICP and BML: writing - original draft; all authors: writing - review & editing; GDP, and JVM: funding acquisition.

## Conflict of interest

The authors declare that they have no conflicts of interest.

## Funding

This work was supported by a sub-award from the University of Illinois to GDP and JVM as part of the research project Realizing Increased Photosynthetic Efficiency (RIPE) that is funded by the Bill & Melinda Gates Foundation, Foundation for Food and Agriculture Research, and the UK Government’s Department for International Development under Grant Number OPP1172157 (funding ended in September 2022).

## Data availability

All relevant data and plant materials are available from the authors upon request. Raw data corresponding to the figures and results described in this manuscript are available online at https://doi.org/10.17632/vncj8cn6xs.1 (Rottet *et al*., 2024). Additional data reported in this paper are presented in the Supplementary Data.

## Abbreviations

ABC: ATP-binding cassette
*A*/*Ci*: CO_2_ assimilation rate as a function of intercellular CO_2_
At: Arabidopsis thaliana
BCT1: bicarbonate transporter 1
CA: carbonic anhydrase
CA-free: specialized *E. coli* strain that lacks Cas
CCM: CO_2_-concentrating mechanism
Ci: inorganic carbon
*cmp*: cytoplasmic membrane protein
cTP: chloroplast transit peptide
IEM: inner envelope membrane
IMAC: immobilized metal affinity chromatography
IMS: intermembrane space
IPTG: isopropyl β-D-1-thiogalactopyranoside
LB: lysogeny broth
NBD: nucleotide-binding domain
OEM: outer envelope membrane
PCR: polymerase chain reaction
Ps: Pisum sativum
SBP: substrate-binding protein
SP: signal peptide
TMD: transmembrane domain
WT: wild-type

## Notes

### Competing Interest Statement

The authors have declared no competing interest.

### Summary of Updates

Supplementary data files added.

