## Supplementary Figures S1-S7+Table S3 for "Engineering the cyanobacterial ATP-driven BCT1 bicarbonate transporter for functional targeting to C_3_ plant chloroplasts"

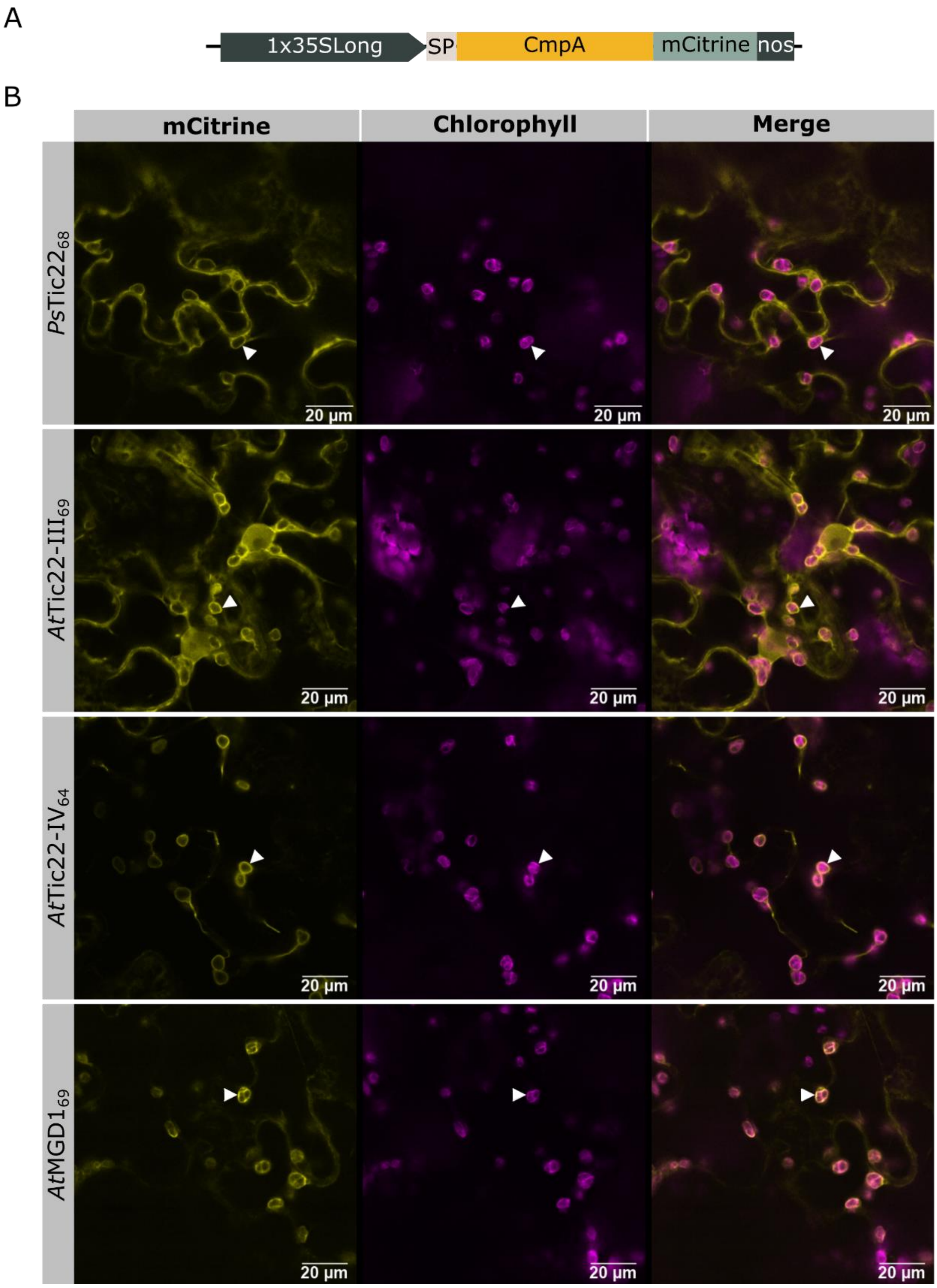


***Figure S1 Targeting of CmpA to the chloroplast intermembrane space.***

**(A)** Schematic of the genetic constructs used in this figure. The chloroplast transit peptides (SP) originate from *Pisum sativum (Ps)* or *Arabidopsis thaliana (At).* The proteins used are *Ps*Tic22 (XP_050885018, GL200), *At*Tic22-III (At3g23710, GL201), *At*Tic22-IV (At4g33350, GL202), and *At*MGD1(At4g31780, GL203). The length of the cTPs are shown as the number of residues in subscript. **(B)** Confocal microscopy images of *N. benthamiana* leaf surfaces transiently expressing CmpA-mCitrine with the various cTPs tested.

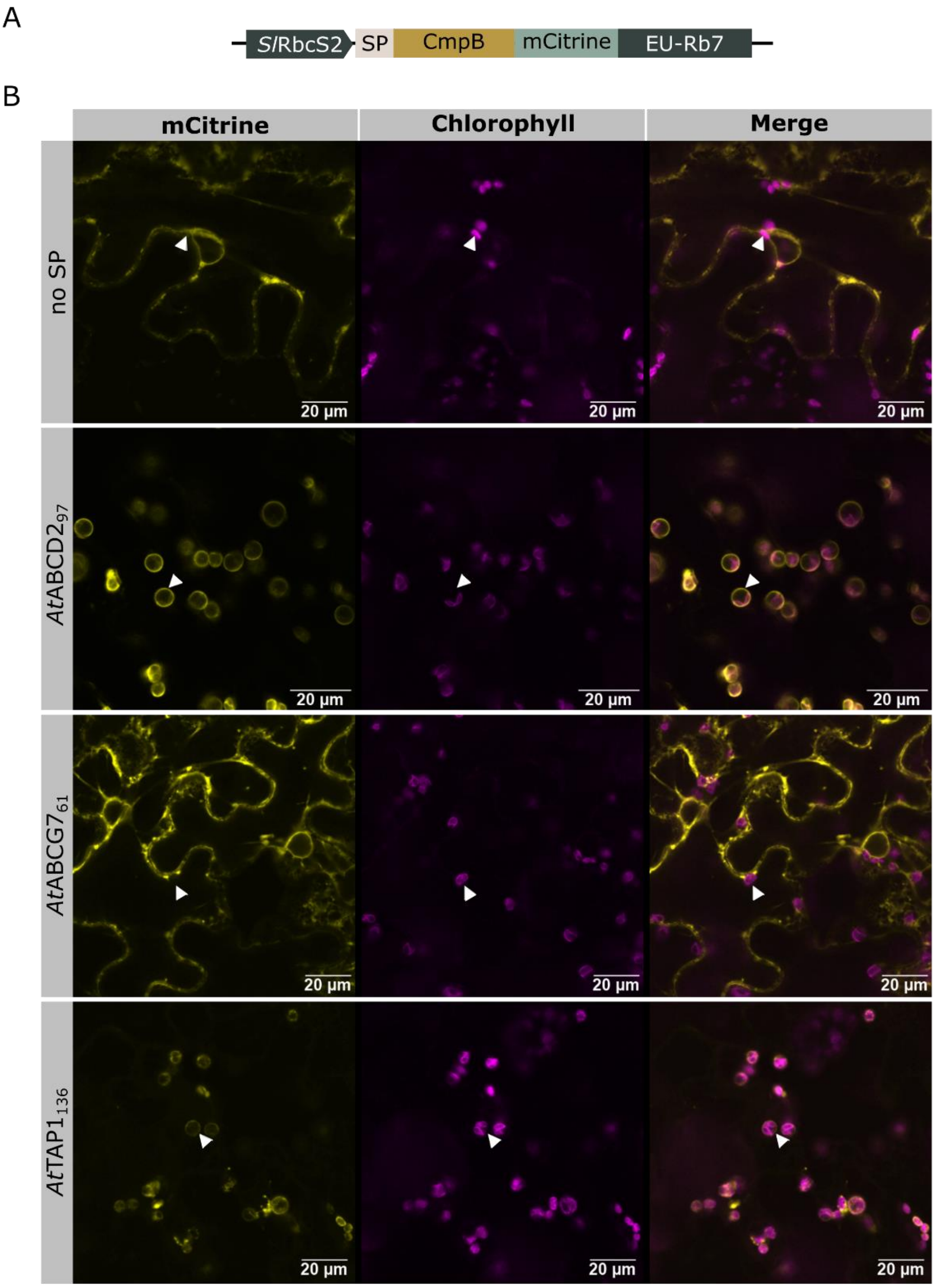


***Figure S2 Targeting of CmpB to the chloroplast inner envelope membrane. (continued)***

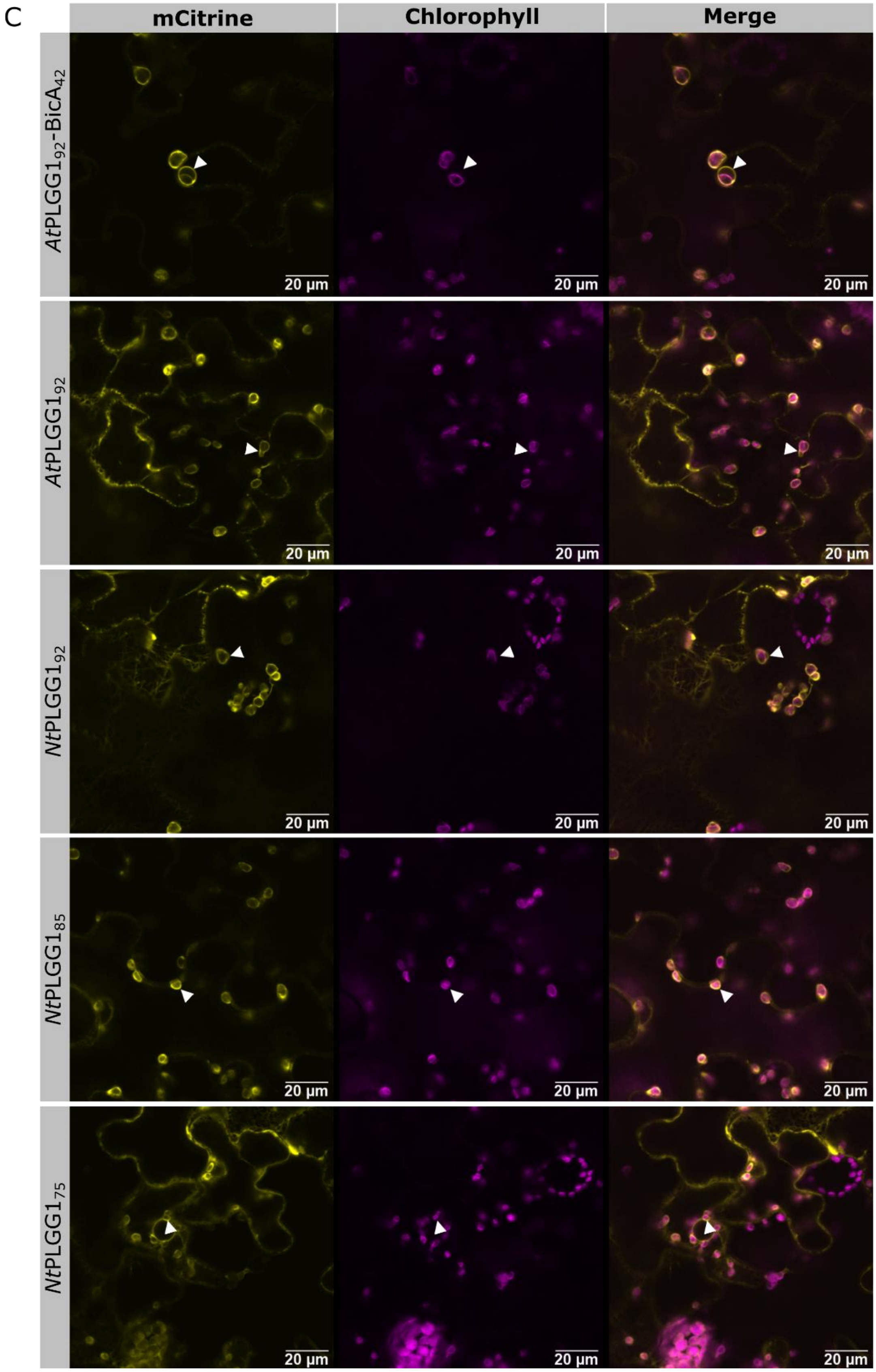


***Figure S2 Targeting of CmpB to the chloroplast inner envelope membrane. (continued)***

***Figure S2 Targeting of CmpB to the chloroplast inner envelope membrane.***

**(A)** Schematic of the genetic constructs used in this figure. The chloroplast transit peptides (SP) originate from *Nicotiana tabacum* (*Nt*), *Arabidopsis thaliana* (*At*) or *Synechococcus sp.* PCC7002. The leader sequences of the following proteins were used: *At*ABCD2 (At1g54350, GL273), *At*ABCG7 (At2g01320, GL274), *At*TAP1 (At1g70610, GL275), *At*PLGG1 (At1g32080, GL267-268), *Nt*PLGG1 (XP_016509146, GL269-271), and BicA (SYNPCC7002_A2371, GL267). The length of the cTPs are shown as the number of residues in subscript. **(B-C)** Confocal microscopy images of *N. benthamiana* leaf surfaces transiently expressing CmpB-mCitrine with the various cTPs tested.

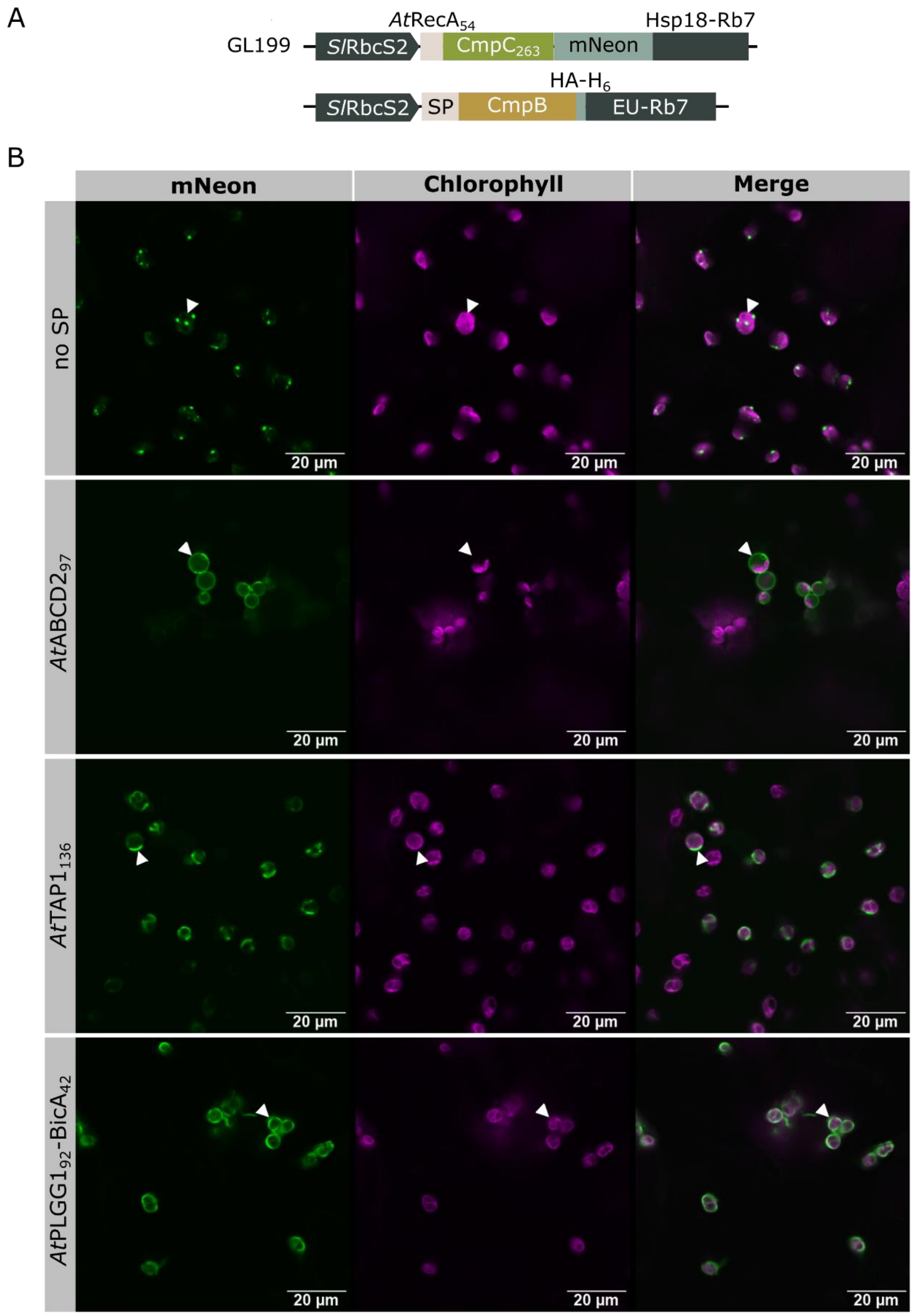


***Figure S3  Orientation of CmpB in the inner envelope membrane. (continued)***

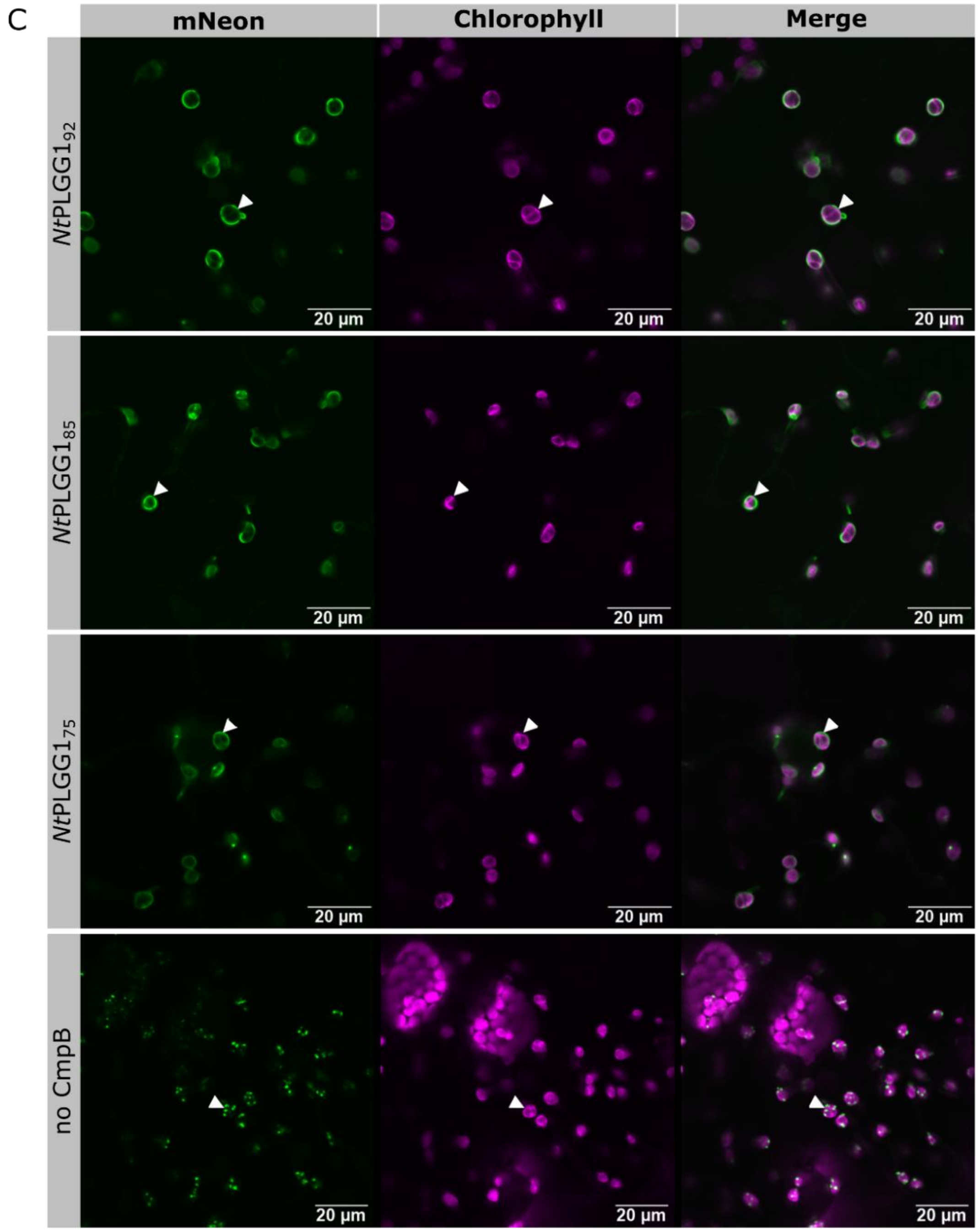


***Figure S3 Orientation of CmpB in the inner envelope membrane.***

**(A)** Schematic of the genetic constructs used in this figure. The chloroplast transit peptides (SP) originate from *Nicotiana tabacum* (*Nt*), *Arabidopsis thaliana* (*At*) or *Synechococcus sp.* PCC7002. The leader sequences of the following proteins were used: *At*ABCD2 (At1g54350, GL239), *At*TAP1 (At1g70610, GL241), *At*PLGG1 (At1g32080, GL244), *Nt*PLGG1 (XP_016509146, GL235-237), and BicA (SYNPCC7002_A2371, GL244). The length of the cTPs are shown as the number of residues in subscript. **(B-C)** Confocal microscopy images of *N. benthamiana* leaf surfaces transiently co-expressing $CmpC_{263}$-mNeon with CmpB-$HAH_6$ with the various cTPs tested.

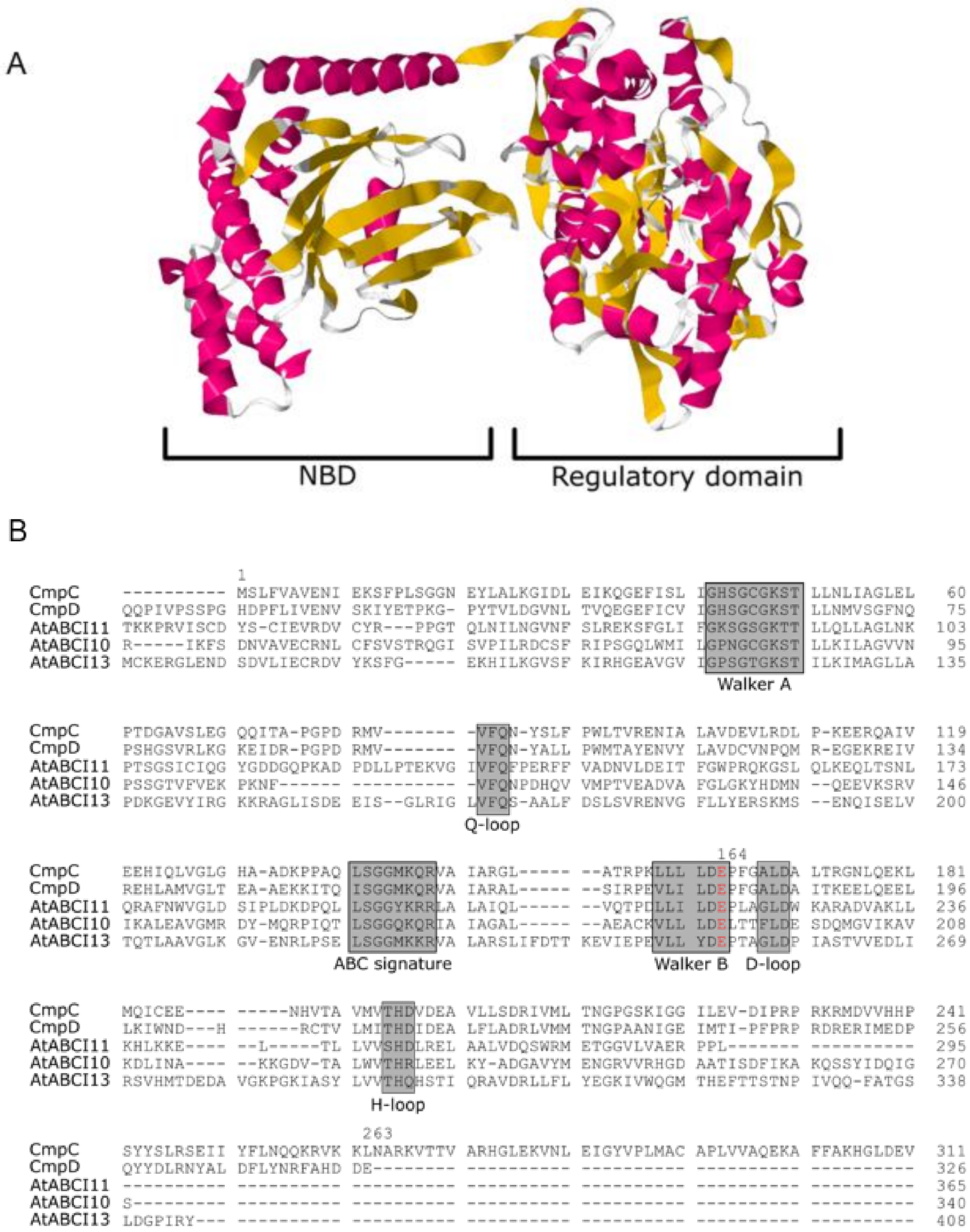


***Figure S4 Structure of CmpC.***

**(A)** 3D structure of CmpC from *Synechococcus elongatus* PCC7942 predicted with RoseTTAFold (Baek *et al.*, 2021) and visualized with Geneious Prime software. CmpC is a 663-residue protein, of which only 263 residues fold in a nucleotide-binding domain (NBD) typical of ABC transporters. The remaining 400 residues fold in a regulatory domain that resembles CmpA. **(B)** Clustal alignment of the highly conserved NBD of CmpC (YP_400507) with CmpD (YP_400508), and other NBD proteins from *Arabidopsis thaliana* (ABCI10, At4g33460; ABCI11, At5g14100; ABCI13, At1g65410). Only the first 311 residues of CmpC are shown.

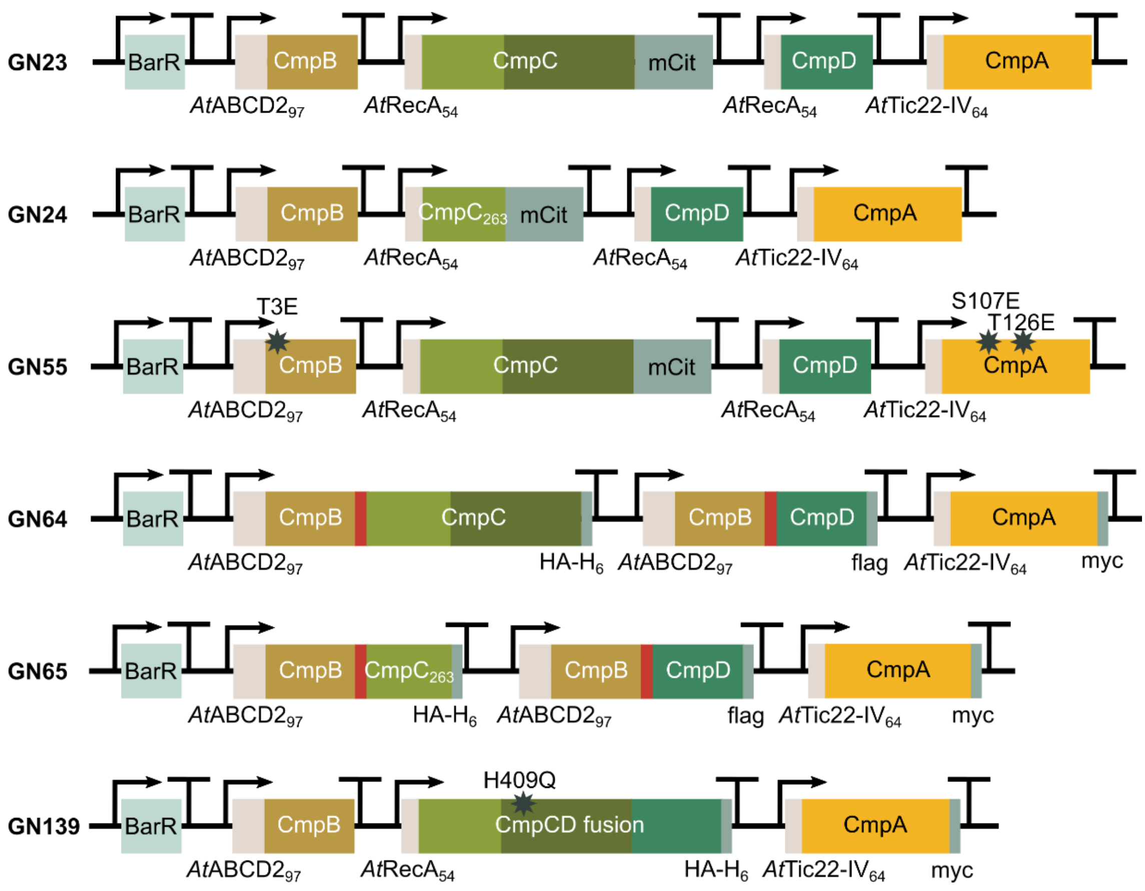


***Figure S5 Genetic constructs screened in Arabidopsis βca5 mutant.***

In the *βca5* mutant, the absence of CA activity in root plastids leads to a very poor growth phenotype at ambient $CO_2$ concentrations (Weerasooriya *et al.*, 2022). This phenotype was previously complemented by expressing a $HCO_3^-$ transporter in root plastids (Förster *et al.*, 2023). Six genetic BCT1 constructs were specifically designed for expression in the Arabidopsis *βca5* mutant. For instance, promoters such as *At*Ubi10 and 35S were used to ensure expression in root plastids (see *Table S2* for details). For selection purposes, a constitutively expressed BASTA resistance (BarR) cassette was included at the start of the constructs. BCT1 proteins were targeted to plastids using targeting sequences as described in *Figure 2*. Short description of the constructs are as follows: unmodified (GN23), no regulatory domain (GN24), phosphorylation mimic (GN55), half-transporter (GN64), half-transporter with $CmpC_{263}$ (GN65), and CmpCD fusion (GN139).

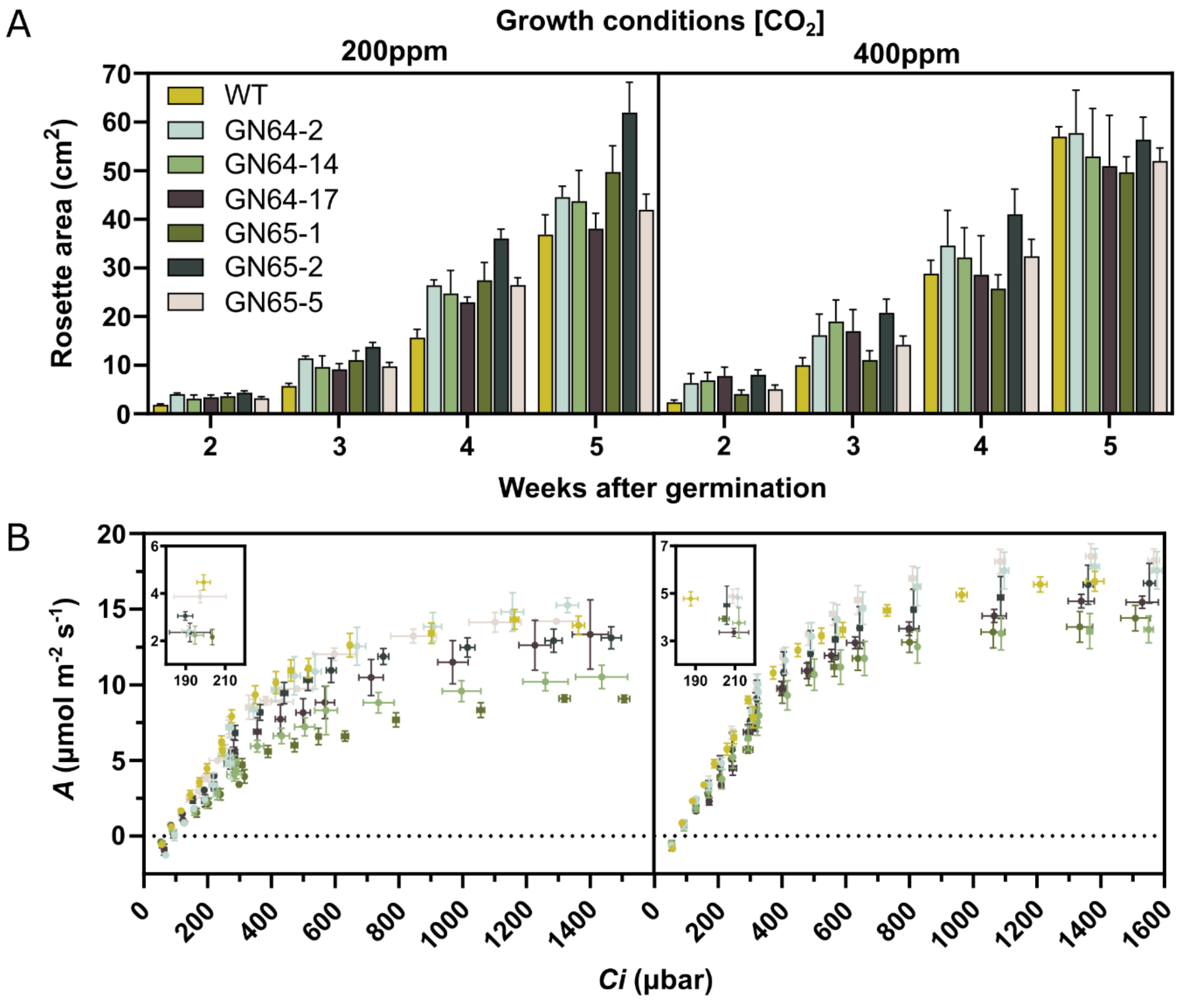


***Figure S6 Rosette area and assimilation rate in transgenic Arabidopsis.***

Two BCT1 mutants were transformed in *A. thaliana* Col-0 background (GN64 and GN65; see *Figure 8*). Four plants for each of the three independent lines were grown at ambient (400 ppm) or low $CO_2$ concentrations (200 ppm). **(A)** Overhead images of plants were taken weekly, and rosette areas were measured as pixel area using the PhenoImage and ImageJ software. Mean ±SE (*n*=4). **(B)** $CO_2$ response curves for Assimilation (*A*; μmol $m^{-2}$ $s^{-1}$) in response to intercellular $CO_2$ (*Ci*; μbar) curves were generated from 50 to 1700 μbar $CO_2$. The inset is a magnified view surrounding 200 μbar (Ci). Mean ±SE (*n*=4, 3 or 2; see *Table S3*).

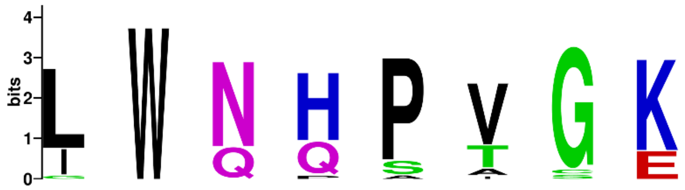


| | | | 288 | 319 | 367 | 409 | 435 | 452 | 453 | 482 |
|---|---|---|---|---|---|---|---|---|---|---|
| CmpC | β | *S. elongatus* PCC7942 | L | W | N | H | P | I | G | K |
| | | *Synechocystis* PCC6803 | L | W | N | H | P | V | S | K |
| | | *Nostoc* PCC7120 | L | W | N | H | P | V | G | K |
| | | *Cyanothece* PCC7425 | L | W | N | H | P | V | G | K |
| | | *T. elongatus* BP-1 | L | W | N | H | P | V | G | K |
| | α | *Synechococcus* WH5701 | L | W | N | H | P | V | G | K |
| CmpA | β | *S. elongatus* PCC7942 | I | W | N | Q | S | T | G | E |
| | | *Synechocystis* PCC6803 | I | W* | N* | Q* | P | T | G | E* |
| | | *Nostoc* PCC7120 | I | W | N | Q | S | T | G | E |
| | | *Cyanothece* PCC7425 | I | W | N | Q | A | T | G | E |
| | | *T. elongatus* BP-1 | I | W | N | Q | P | T | G | E |
| | α | *Synechococcus* WH5701 | I | W | N | Q | S | T | G | E |
| NrtA | β | *S. elongatus* PCC7942 | L | W | Q | H | P | V | G | K |
| | | *Synechocystis* PCC6803 | L* | W* | Q* | H* | P* | V* | G* | K* |
| | | *Nostoc* PCC7120 | L | W | Q | H | P | V | G | K |
| | | *Cyanothece* PCC7425 | L | W | Q | H | P | V | C | K |
| | | *T. elongatus* BP-1 | L | W | Q | H | P | V | G | K |
| | α | *Synechococcus* WH5701 | L | W | Q | H | P | V | G | K |
| NrtC | β | *S. elongatus* PCC7942 | G | W | G | P | P | A | G | K |
| | | *Synechocystis* PCC6803 | L | W | N | Q | P | V | G | K |
| | | *Nostoc* PCC7120 | L | W | N | Q | P | A | G | K |
| | | *Cyanothece* PCC7425 | L | W | N | H | P | V | G | K |
| | | *T. elongatus* BP-1 | L | W | N | H | P | V | G | K |
| | α | *Synechococcus* WH5701 | L | W | N | H | P | V | G | K |

***Figure S7 Sequence alignment and corresponding WebLogo conservation sequence of CmpC, CmpA, NrtA and NrtC from β- and α-cyanobacteria.***

Amino acids marked with an asterisk were identified by Koropatkin et al., (2006) as putative ligand binding residues. Residues in NrtA from *Synechocystis* PCC6803, responsible for the transport of nitrate were highly conserved between species and could be mapped onto other proteins from different species. Here, we identified CmpC residue H409 as the equivalent of the putative ligand-binding residue H196 in NrtA 6803 (Koropatkin *et al.*, 2006) and Q198 in CmpA 6803 (Koropatkin *et al.*, 2007). Key changes in putative ligand-binding residues were postulated to be responsible for the difference in substrate specificity between NrtA (nitrate) and CmpA (bicarbonate). Sequence alignment were performed using the Clustal Omega algorithm (Geneious Prime 2022.1.1). Displayed residue numbers are nonsequential and have been numbered according to the *S. elongatus* PCC7942 CmpC sequence. Polar amino acids are coloured green, basic amino acids are blue, acidic amino acids are red and hydrophobic amino acids have been coloured black.

### *Table S1 List of primers used in this study*

This Excel spreadsheet contains the list of primers used in this study. The first part of Table S1 is dedicated to cloning and domestication (Dom) primers or oligos used to generate CDS2ns parts (Engler *et al.*, 2014). The rest of Table S1 lists additional primers that were used for colony PCR and sequencing.

### *Table S2 List of constructs used in this study*

This Excel spreadsheet contains the list of genetic constructs designed and used in this study. The constructs are grouped in separate tabs according to their construct level and target organism. **Tab 1** lists all level 1 (GL) constructs used for transient expression in *N. benthamiana*, **tab 2** lists all level 2 (GM) constructs used for expression in *E. coli.*, **tab 3** lists the level 3 (GN) constructs used for expression in *E. coli.*, and **tab 4** lists the six level 3 (GN) constructs used to stably transform *A. thaliana*.

The various parts types shown in the top rows were previously described for Golden Gate Modular Cloning (Engler *et al.*, 2014). Pro, promoter; 5U, 5′ untranslated sequence; 5U(f), 5′ untranslated sequence for N terminal fusions; NT, N terminal tag or localization signal; SP, signal peptide; CDS, coding sequence; CT, C terminal tag or localization signal; 3U, 3′ untranslated sequence; Ter, terminator; ns, no stop codon.

The acceptor plasmids, pOdd1-4 and pEven1-4, were previously described for Loop Assembly (Pollak *et al.*, 2019). The terminal acceptor plasmids, pFA-Odd and pFA-Even, were designed and assembled in-house to be compatible with Loop Assembly (Pollak *et al.*, 2019). In brief, the pFA31 backbone (Flamholz *et al.*, 2020) was modified by changing its antibiotic resistance, introducing compatible cloning sites and replacing TetR with the LacIQ-pTrc-pLac repressor/promoter cassette.

The short Odd and Even dummies were also designed in-house using a random sequence generator http://www.faculty.ucr.edu/~mmaduro/random.htm and the appropriate 4 bp overhangs for each position(s) the dummies were used to fill in. These were then either ordered from Macrogen Inc. (Seoul, South Korea) as primers and annealed in-house, or ordered as double-stranded oligos.

***Table S3***

**Table S3**. $CO_2$ compensation points for BCT1 transformants in WT Arabidopsis.

| Growth [$CO_2$] | Genotype | *n* | $CO_2$ compensation point (µbar) |
|---|---|---|---|
| 200 ppm | WT | 4 | 70.95 ± 1.95 |
| | GN64-2 | 3 | 101.10 ± 1.00 |
| | GN64-14 | 3 | 79.59 ± 6.35 |
| | GN64-17 | 3 | 89.31 ± 6.80 |
| | GN65-1 | 3 | 79.72 ± 3.55 |
| | GN65-2 | 4 | 70.46 ± 5.46 |
| | GN65-5 | 2 | 69.23 ± 5.15 |
| 400 ppm | WT | 4 | 71.11 ± 2.07 |
| | GN64-2 | 4 | 70.31 ± 4.67 |
| | GN64-14 | 4 | 68.89 ± 2.74 |
| | GN64-17 | 3 | 70.53 ± 1.55 |
| | GN65-1 | 3 | 69.56 ± 6.81 |
| | GN65-2 | 4 | 71.68 ± 6.21 |
| | GN65-5 | 4 | 70.90 ± 2.79 |

The number of replicates is indicated (*n*). Mean ± SE.
